# Programmable DNA-peptide nanostructures for multivalent regulation of intracellular signalling

**DOI:** 10.64898/2026.09.03.749096

**Authors:** Maria Zacharopoulou, Zoya Cassidy, Siding Qin, Zixuan Huo, Sergio Grannum, Aditya Sridhar, Joseph E. Chambers, Marc de la Roche, Laura S. Itzhaki, Ioanna Mela

**Affiliations:** Department of Pharmacology, University of Cambridge, Tennis Court Road, Cambridge CB2 1PD, UK; Cambridge Institute for Medical Research, Hills Road, Cambridge CB2 0XY, UK; Department of Biochemistry, University of Cambridge, Tennis Court Road, Cambridge CB2 1GA, UK

**Author notes:** Centre for Organismal Studies (COS), Heidelberg University, Im Neuenheimer Feld 230, cS120 Heidelberg, Germany.

## Abstract

The Wnt/β-catenin pathway is constitutively active in most colorectal and other cancers. Tankyrase (TNKS) promotes Wnt signalling by PARylating AXIN, a rate-limiting scaffold subunit of the β-catenin destruction complex, and its inhibition is a validated strategy for pathway downregulation. Existing small-molecule TNKS inhibitors that target the catalytic PARP domain suffer from off-target effects across the PARP family. Here, we present an alternative approach that targets the substrate-binding domains of TNKS using a short TNKS-binding peptide (TBP) presented multivalently on DNA nanostructures. Such a strategy may be required to effectively disrupt the function of targets such as TNKS, which is known to form high-order assemblies in the cell. We engineered DNA nanostructures with two distinct geometries: a 2D triangle (∼100 nm) displaying 27 copies of TBP, and a compact 3D tetrahedron (∼10 nm) displaying 2 TBP copies. We show that both nanostructures assemble efficiently, can be functionalised with TBP in high yields, and retain binding to TNKS protein *in vitro*. DNA nanostructures are efficiently internalised to the cytoplasm by HeLa Kyoto and colorectal cancer cells, with TBP functionalisation enhancing rather than hindering uptake. In HeLa cells, both triangle-TBP and tetrahedron-TBP downregulated Wnt signalling to a similar extent despite an order-of-magnitude difference in TBP copy number, while the same concentration of free TBP had no effect. This finding likely reflects the two functions of the DNA nanostructures - intracellular delivery and multivalent display - whereby the smaller nanostructures more efficiently internalise the TBP ligand but have lower valency. These results establish DNA nanostructures as a modular, tuneable platform for multivalent inhibition of intracellular clustered targets such as TNKS.

## Introduction

The Wnt signalling pathway is evolutionarily conserved and plays a major role in tissue homeostasis, regeneration, and embryogenesis. Dysregulation of the Wnt signalling pathway has been heavily implicated in cancer initiation, progression, metastasis, and resistance to treatment [1]. In colorectal cancer, Wnt pathway mutations occur in approximately 93% of cases, with ∼81% of non-hypermutated tumours exhibiting constitutive pathway activation [2]. This hyperactivation arises from the destabilisation of the β-catenin destruction complex, a multiprotein assembly which consists of the scaffold proteins AXIN and adenomatous polyposis coli (APC), and two kinases that phosphorylate β-catenin (GSK3β and CK1α) (Figure 1A). A key regulator of the Wnt pathway is TNKS. TNKS-mediated PARylation of AXIN is necessary for its degradation [3], [4]. TNKS refers to the proteins tankyrase-1 and its paralog tankyrase-2, which share a highly conserved catalytic PARP domain and largely overlapping functions in regulating Wnt signalling. AXIN is present at lower levels than other key components of the β-catenin destruction complex, so it is theorised to act as the rate determining step in complex assembly [3]. Thus, inhibiting TNKS and, thereby, AXIN degradation, provides promising therapeutic potential for stabilisation of the β-catenin destruction complex and reduction of downstream Wnt signalling.

**Figure 1:**
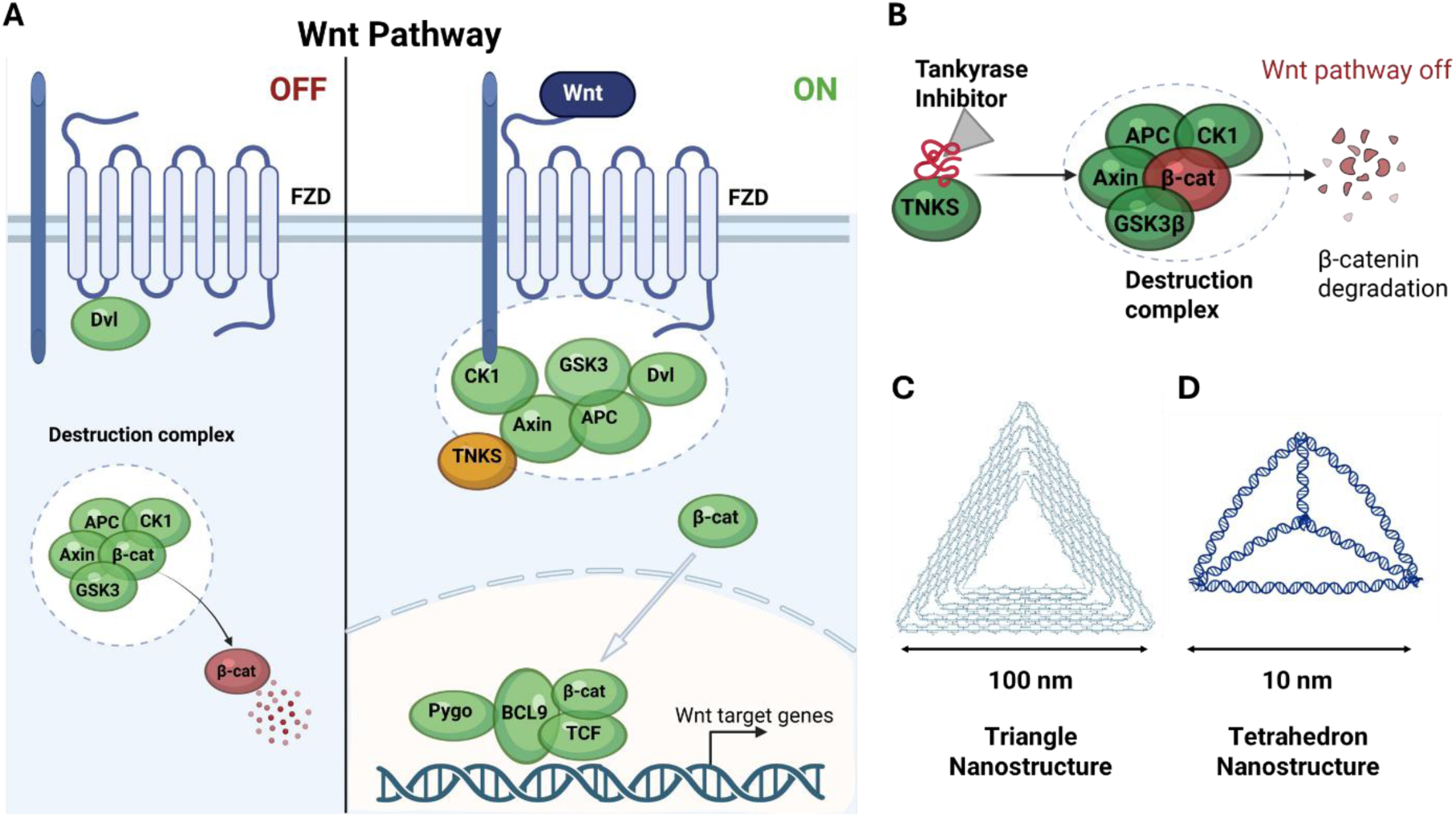
**A. Regulation of Wnt/β-catenin signalling by the destruction complex and TNKS**. Without Wnt ligand, the destruction complex (AXIN, APC, GSK3β, CK1) binds β-catenin, driving its phosphorylation and proteasomal degradation to keep proliferative target genes off. Wnt binding to Frizzled/LRP5/6 inactivates the destruction complex, allowing β-catenin to accumulate, enter the nucleus, and activate TCF/LEF-driven proliferation genes. Tankyrase promotes this “on” state by PARylating AXIN, marking it for degradation and thereby destabilising the destruction complex. Targeting TNKS for inhibition is a promising strategy to downregulate the Wnt pathway. **B. DNA nanostructures decorated with TNKS binding peptides (TBP) inhibit TNKS,** thereby stabilising AXIN and the destruction complex, leading to β-catenin degradation and downregulation of the Wnt pathway**. C. Triangle nanostructure** (2D, 100 nm), also referred to in the literature as the “Rothemund triangle”, **D. Tetrahedron nanostructure** (3D, 10 nm), comprising 4 DNA strands. Created in Biorender.com.

TNKS dysregulation has been linked to a variety of cancers including colorectal, breast, fibrosarcoma, ovarian, glioblastoma, pancreatic, and gastric cancer highlighting TNKS inhibition as a broadly relevant therapeutic strategy [5]. A number of studies have reported the development of small molecule inhibitors of the TNKS catalytic activity as a means of modulating Wnt pathway activity [6], [7], [8], [9], [10]. However, these small molecule inhibitors target the catalytic PARP domain and may lack specificity for TNKS over other PARP family members [11], [12], leading to off-target effects. Indeed, gastrointestinal off-target toxicity was observed in mouse models, which limits their therapeutic application [13]. Moreover, PARP-domain inhibitors do not block TNKS’s non-catalytic scaffolding functions mediated by its ankyrin repeat clusters (ARCs). Targeting the ARC domains therefore offers a route to inhibit both catalytic and non-catalytic TNKS functions with improved specificity.

A consensus tankyrase-binding peptide (TBP) motif recognised by the ARC domains has been identified, and we previously demonstrated that macrocyclised TBPs bind TNKS and inhibit Wnt signalling. [14] [15]. We further showed that grafting TBP onto the inter-repeat loops of a consensus-designed tetratricopeptide repeat protein (CTPR) scaffold generates multivalent binding proteins that cluster TNKS intracellularly and enhance pathway inhibition [16]. In each case, delivery of the peptide/protein into the cell required further functionalisation, either using a cell-penetrating peptide motif or by lipid nanoparticle encapsulation.

Indeed, multivalency is an important parameter in targeting TNKS, as increasing evidence points to TNKS itself being a multivalent target. TNKS self-assemble into higher-order complexes through C-terminal SAM domain polymerisation and additionally each TNKS monomer contains multiple N-terminal Ankyrin Repeat Clusters (ARCs) that recruit the substrate [17], [18], [19], [20]. SAM-mediated head-to-tail assembly clusters the catalytic PARP domains, which is essential for allosteric activation and enzymatic PARylation [17]. These multivalent SAM and ARC interactions have also been shown to facilitate liquid–liquid phase separation (LLPS), driving the concentration of tankyrase and key scaffolds such as AXIN into dynamic biomolecular condensates and signalosomes [17], [19], [20], [21]. Within this multimeric architecture, a multivalent targeting approach offers a distinct thermodynamic advantage by exploiting avidity to achieve significantly higher apparent binding affinity and effectively engage clustered TNKS assemblies. Our strategy here is to present multiple copies of TBP on DNA nanostructures (DNA origami), which functions as a multivalent drug delivery vehicle with exceptional programmability.

DNA origami offers an attractive platform for multivalent intracellular targeting. These structures are formed by folding a long single-stranded DNA scaffold with short staple strands into precise two- or three-dimensional geometries [22], or by assembling smaller motifs such as DNA tetrahedra without a scaffold [23]. DNA nanostructures are advantageous over many synthetic drug delivery systems as they are intrinsically biodegradable and have shown internalisation into mammalian cells [24], [25], [26], [27]. They can maintain structural integrity within cells for 24 hours, depending on their physical and chemical properties, and are susceptible to eventual degradation by DNase I present intracellularly and in bodily fluids [28], [29]. Due to their highly customisable and predictable geometries, these DNA nanostructures can serve as targeted drug delivery vehicles by precisely carrying therapeutic payloads—such as small molecules, peptides, nanobodies—in a stoichiometrically controlled manner and with programmable distances between payloads [30], [31].

The internalisation of DNA nanostructures by mammalian cells is an active area of research, particularly because these programmable architectures hold great potential for targeted drug delivery. DNA as a molecule is highly negatively charged and hydrophilic and therefore cannot diffuse across the hydrophobic plasma membrane. Instead, mammalian cells internalise DNA origami primarily via endocytosis [32]. For untargeted or unmodified DNA origami, research indicates that scavenger receptors on the mammalian cell membrane play a primary role in recognising and capturing the negatively charged DNA structures from the extracellular environment [32]. Internalisation efficiency is heavily dependent on nanostructure geometry, size, and compactness [33]. Unmodified DNA nanostructures can be internalised via clathrin and caveolin-mediated endocytosis [32], [34]. The clathrin-mediated pathway facilitates endocytosis of cargo with a diameter <200 nm, whereas nanoparticles of smaller sizes (20–100 nm) can be endocytosed through a caveolae-dependent mechanism. Simply changing the size or shape of a DNA origami structure can drastically alter its cellular uptake pathway and total internalisation efficiency [25], [33], [34]. Other strategies to enhance endocytosis or alter the subcellular localisation on nanostructures rely on using a targeting molecule (such as a nanobody, peptide, aptamer) to bind a specific membrane receptor for receptor-mediated endocytosis [35], [36]. Triangle-shaped DNA origami has been shown to passively accumulate at tumour regions *in vivo* [37], and nanostructures have also been specifically targeted to cell-surface associated tumour markers through attached moieties, such as aptamers and antibodies [37], [38].

A central requirement for therapeutic efficacy in this system is the accessibility of cytosolic TNKS. DNA nanostructures enter mammalian cells primarily through endocytic pathways [39], [40] and therefore targeting cytosolic machinery like the β-catenin destruction complex requires internalised nanostructures to clear the endosome before lysosomal degradation occurs. Emerging evidence indicates that a subpopulation of endocytosed DNA nanostructures can undergo endosomal escape or transient membrane perturbation during endolysosomal maturation, permitting access to the cytoplasm [41], [42], [43], [44]. This intracellular availability is aided by the dense spatial packing of DNA origami, which confers enhanced resistance to endolysosomal nucleases relative to unstructured nucleic acids [45], [46], [47]. Crucially, by presenting multivalent arrays of TBP on a rigid nanostructure, even a modest cytosolic fraction can achieve a high local concentration of binding sites, leveraging avidity to effectively engage cytosolic TNKS assemblies. Here, we employ DNA nanostructures as multivalent delivery vehicles for TBP to inhibit TNKS and downregulate Wnt signalling. To explore how geometry and valency influence cellular uptake and functional inhibition, we designed two architectures: a 2D Rothemund triangle (triangle) [22] (∼100 nm) displaying 27 TBP copies to maximise avidity, and a compact 3D tetrahedron [23] (∼10 nm) displaying 2 TBP copies to favour efficient endocytosis (Figure 1 C,D). We characterise the nanostructure assembly, TBP functionalisation, TNKS binding, cellular internalisation, and Wnt pathway inhibition in HeLa Kyoto and SW480 colorectal cancer cells. We demonstrate that both geometries are internalised and successfully downregulate the Wnt pathway in HeLa cells, revealing how nanostructure architecture and valence can be tailored to balance cellular entry with target inhibition.

## Results

### Nanostructure design, functionalisation and characterisation

Our strategy for modifying the DNA nanostructures to carry TBP was to extend selected structural staples at predesigned positions, creating single-stranded oligonucleotide overhangs (“sticky ends”). Each overhang was designed with one end complementary to a specific site within the nanostructure and a free end available for hybridisation with a complementary oligonucleotide. For the triangle origami, 13–14 overhangs were incorporated on each face to maximise the probability of TNKS engagement as the structure tumbles in solution. A complementary 25-nt oligonucleotide (“oligo-linker”) to this overhang, carrying a 5′-azide modification, was conjugated to TBP via a strain-promoted azide–alkyne cycloaddition with a dibenzocyclooctyne (DBCO) group on the peptide’s N-terminus (Figure 2A). The TBP peptide additionally carried a PEG2 spacer for conformational flexibility and a Cy5 label for downstream detection (DBCO-(PEG)2-C(Cy5)NREAGDGEE). The resulting oligo–peptide conjugate was then hybridised to the DNA origami at the predesigned overhang positions.

**Figure 2:**
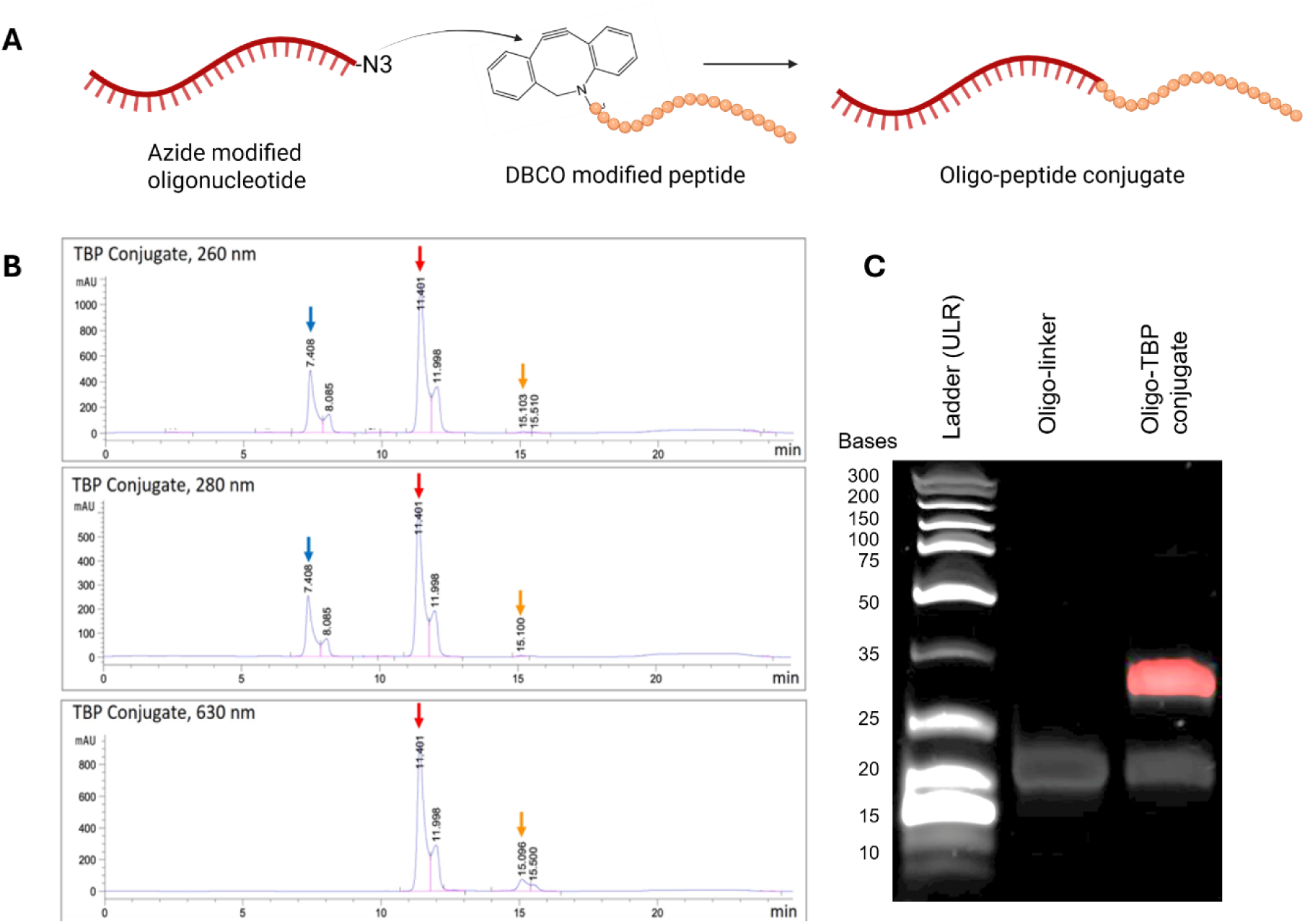
DNA-peptide conjugation and characterisation. **A.** Conjugation strategy. Azide modified oligonucleotide (oligo-linker) is click-reacted to a DBCO modified peptide (TBP) to form an oligopeptide conjugate. This conjugate will then by hybridised to predesigned modification positions on the nanostructures. **B.** HPLC shows effective separation of the click reaction products, while the reaction efficiency is calculated at >85%. Shown are the absorbance at 260 nm, 280 nm, and 630 nm (Cy5). **C.** Acrylamide gels show click reaction product (conjugate) of higher mass (35 bases), which also fluoresces at 647 (Cy5). Unreacted oligonucleotide (linker) can also be detected at the 20 bases mark.

The oligo-linker and TBP were mixed at a 1:1 molar ratio and reacted overnight at 4°C, achieving a conjugation efficiency of >85% as determined by reversed-phase high-performance liquid chromatography (RP-HPLC) (Figure 2B). Three peaks were resolved in the chromatogram, corresponding to the unreacted oligo-linker (7.4 min), the oligo–peptide conjugate (11.4 min), and unreacted TBP (15.1 min). The conjugate peak was collected and used for subsequent hybridisation onto the DNA nanostructures. The click reaction was further validated by denaturing acrylamide gel electrophoresis (Figure 2C), which showed a band at ∼20 bases corresponding to the unreacted DNA linker alongside a higher-molecular-weight band in the conjugate lane. Cy5 fluorescence co-localised with this higher-molecular-weight DNA band, confirming successful formation of the TBP conjugate.

DNA triangle nanostructures were folded as previously established [22] by mixing the structural and modified overhang staples with the scaffold DNA. The TBP conjugate was added to the folded origami in 10-fold molar excess and hybridised to the overhang positions (40 °C, 1 h) (Figure 3A). Nanostructures were purified by centrifugation, and their morphology was assessed by atomic force microscopy (AFM). Because TBP is conjugated via a flexible a DNA overhang and a PEG linker, it is inherently challenging to resolve; we therefore imaged samples in FastScan Tapping Mode, applying the minimal force necessary to visualise the peptide without disrupting its position. The bare triangle nanostructures were well-formed, showing distinct trapezoidal domains (∼100 nm across the outer perimeter) connected by bridging staples (Supplementary Figure S1). Incorporated peptides were detected on the surface of the modified triangles as discrete spots of increased height (∼1.5 nm, Supplementary Figure S2), corresponding well to the designed modification positions on the triangle map (Figure 3A, black dots). An average of 11 modification positions was detected per triangle, fitting with the 13–14 predesigned modifications per triangle plane.

**Figure 3:**
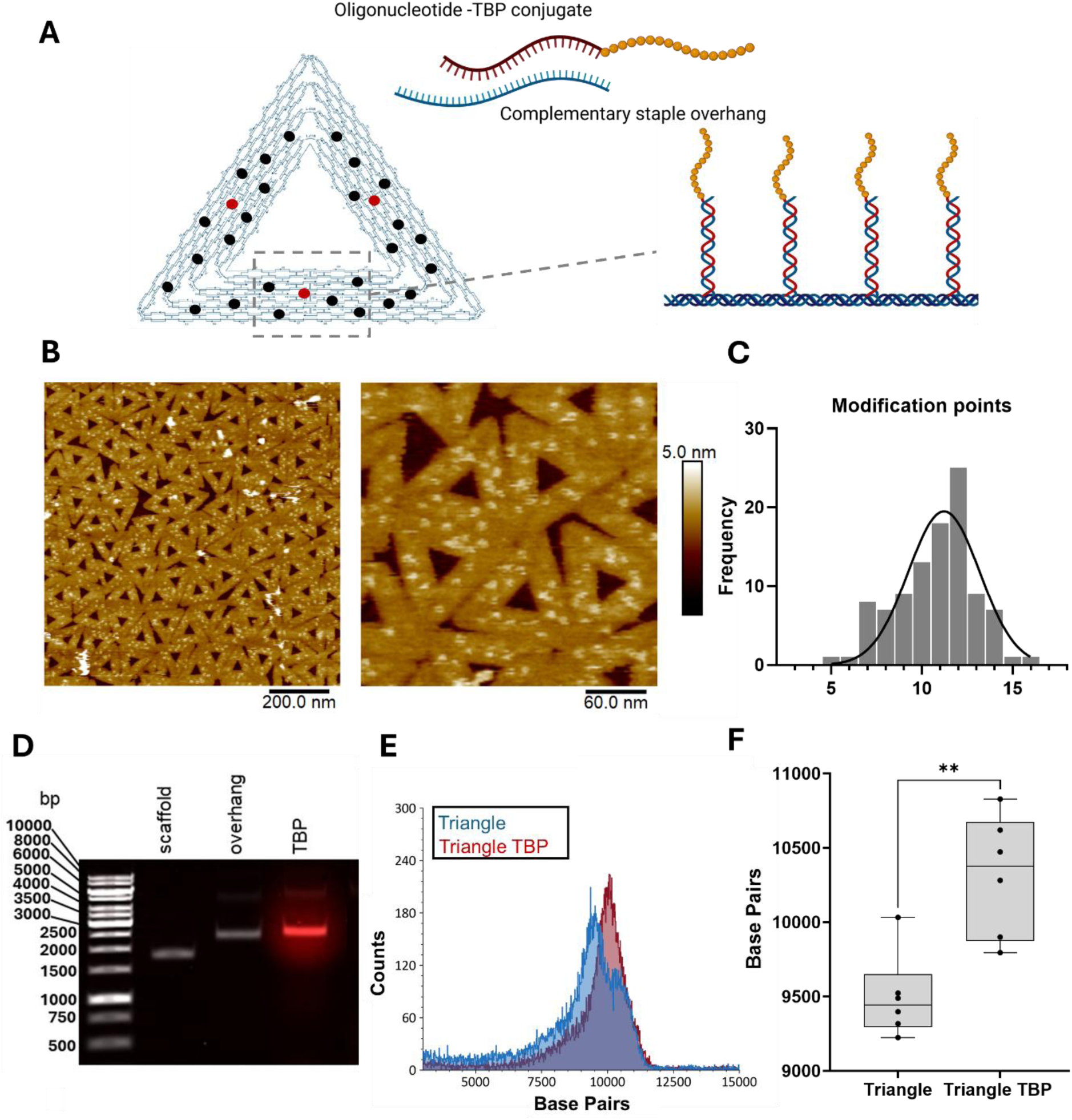
Triangle nanostructure characterisation. **A.** Schematic of the hybridisation of the oligo-peptide conjugate on the triangle planes. The TBP extends from both planes of the triangles in predesigned positions (black dots on triangle map, red dots correspond to fluorophore positions). Created with Biorender.com. **B.** AFM of the hybridisation products shows points of increased height (2 nm) on the hybridisation positions. **C.** Histogram of the triangle modification points detected by AFM. A Gaussian distribution was fitted to the data, with average modifications per triangle 11.2 ± 1.9. **D.** Agarose gels (1%) showing positive shift in mass upon origami folding (lanes 1 vs 2) and further shift in mass (lane 3) upon TBP hybridisation. The band in lane 3 fluoresces, due to presence of Cy5 on TBP. **E.** Mass Photometry distribution of unmodified Triangle and Triangle TBP shows a positive shift in the distribution upon hybridisation. **F.** Mass photometry average of three biological repeats shows significant difference in mass upon TBP hybridisation.

Hybridisation efficiency was further validated by agarose gel electrophoresis (Figure 3D). Comparing the scaffold DNA (lane 1) to the folded triangle nanostructure with overhangs (lane 2) revealed a positive mass shift, reflecting the different migration behaviour of the folded nanostructure relative to unfolded DNA. A further positive shift was observed upon TBP hybridisation, accompanied by overlapping Cy5 fluorescence, confirming successful incorporation of TBP. Mass photometry was also explored as an alternative method for detecting mass changes due to functionalisation; to our knowledge, this is the first application of mass photometry to measure DNA origami mass. Incorporating 20 mM MgCl₂ into the origami buffer yielded sufficient landing events to resolve both unmodified and TBP-functionalised nanostructures (Figure 3E). A significant increase in mass was observed for the triangle upon TBP hybridisation (+820 bp) (Figure 3F), further corroborating TBP hybridisation onto the triangle planes. It must be noted that mass calibration was performed using low range MW DNA ladder, which is linear DNA and thus expected to result in different landing events compared to folded DNA origami; for that reason the absolute bp values are not as expected (triangle origami should measure at ∼7200 bp in contrast to ∼9000 bp), and we expected a shift in mass of approximately +100 bp upon addition of 27 TBP copies (each TBP weighs 2.2 kDa), rather than +820 bp - however the relative differences between samples are still valuable.

To modify the tetrahedron with TBP, we extended two vertices with the same overhang sequence used for the triangle origami, enabling downstream hybridisation with the TBP conjugate (Figure 4A). We first tested whether the tetrahedron could fold stably with these two overhangs. Agarose gels of different strand combinations (Figure 4B); showed that only the complete four-strand mixture produced the characteristic tetrahedral band (lane 7), indicating efficient folding as previously established [23] and substitution of strands 1 and/or 4 with their overhang-bearing counterparts (lanes 8–10) yielded correctly folded structures with the expected mass shifts.

**Figure 4:**
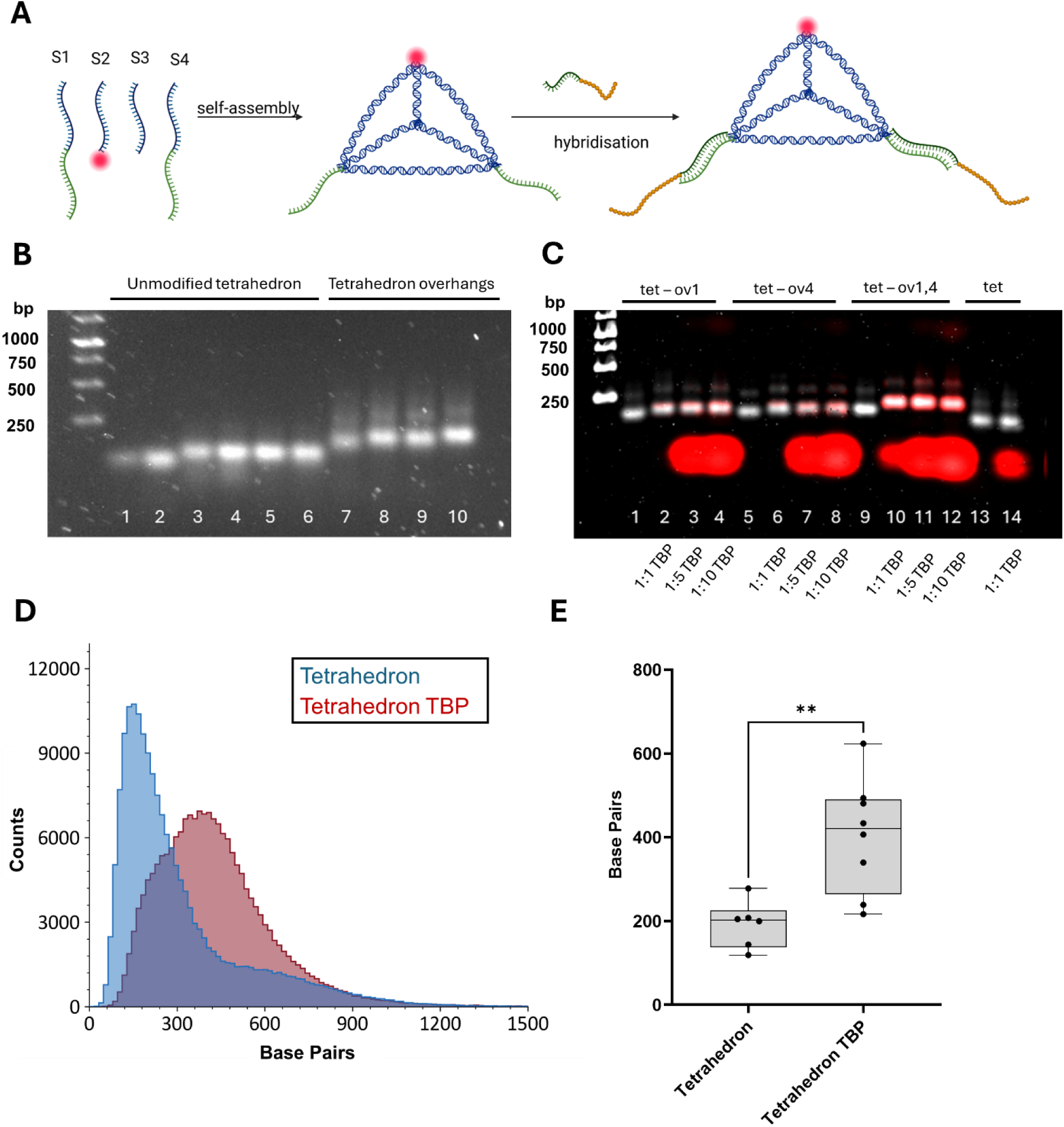
Tetrahedron nanostructure characterisation. **A.** Schematic of the hybridisation of the oligo-peptide conjugate on the tetrahedron vertices. Two of the vertices are extended with the same overhang sequence as used for the triangle and hybridised to the TBP conjugate. **B.** Agarose gels showing efficient formation of the tetrahedron, when modified with two overhangs. Lanes are as follows: 1: strand1 + strand2, 2: strand3 + strand4, 3: strand1 + strand2 + strand3, 4: strand1 + strand2 + strand4, 5: strand1 + strand3 + strand4, 6: strand2 + strand3 + strand4, 7: strand1 + strand2 + strand3 + strand4, 8: strand 1 overhang and strands 2-4, 9: strands 1-3 and strand4 overhang, 10: strand 1 overhang, strand 2, strand 3, strand 4 overhang. **C.** Agarose gels showing effective hybridisation of oligo-peptide conjugates on the tetrahedra. Lanes are as follows: 1: tetrahedron with strand1 overhang, 2: tetrahedron with strand1 overhang + 1:1 TBP, 3: tetrahedron with strand1 overhang + 1:5 TBP, 4: tetrahedron with strand1 overhang + 1:10 TBP, 5: tetrahedron with strand4 overhang, 6: tetrahedron with strand4 overhang + 1:1 TBP, 7: tetrahedron with strand4 overhang + 1:5 TBP, 8: tetrahedron with strand4 overhang + 1:10 TBP, 9: tetrahedron with strand1 and strand4 overhang, 10: tetrahedron with strand1and strand4 overhang + 1:1 TBP,11: tetrahedron with strand1 and strand4 overhang + 1:5 TBP, 12: tetrahedron with strand1 and strand4 overhang + 1:10 TBP, 13: tetrahedron without overhang, 14: tetrahedron without overhang + 1:1 TBP**D.** Mass photometry of the bare and TBP-modified tetrahedron show positive shift in mass upon hybridisation. **E.** An average of three biological repeats of mass photometry show significant change in mass upon hybridisation corresponding to two TBP copies.

We next tested whether the overhang positions were accessible for hybridisation with the TBP conjugate (Figure 4C). Tetrahedra bearing an overhang at strand 1 were incubated with increasing concentrations of TBP (molar ratios 1:1, 1:5, and 1:10; lanes 1–4), and the same was performed for the strand 4 overhang variant (lanes 5–8) and the double-overhang variant (lanes 9–12). A shift in migration was detected at a 1:1 overhang:TBP ratio, which did not increase further at higher TBP ratios. Cy5 fluorescence confirmed TBP incorporation into the tetrahedra bands, with significant free, unhybridised peptide detected beneath the tetrahedra bands at 1:5 and 1:10 ratios; no significant free peptide was detected at the 1:1 ratio, indicating efficient hybridisation. Unmodified tetrahedra (lanes 13 and 14) migrated at lower molecular weight, showed no shift upon incubation with TBP, and displayed Cy5 fluorescence only in the free-peptide position (lane 14). Finally, mass photometry detected the DNA tetrahedron at approximately the expected molecular weight (200 bp, Figure 4D) and showed a significant increase in mass upon TBP hybridisation, (Figure 4D–E), although it must be noted that the increase in mass is again larger than expected, as described above.

To confirm that TBP conjugated to the DNA nanostructures retains its ability to bind the target, we tested *in vitro* binding to a TNKS fragment (residues 488–649, ARC4 domain, containing a single TBP-binding site). Triangle nanostructures incubated with TNKS-ARC4 (1:10 ratio, RT, 30 min), purified by filter centrifugation, and imaged by AFM showed spherical features of 2–3 nm on the triangle surface, consistent in size with TNKS-ARC4, a 25 kDa protein (Figure 5A, Supplementary Figure S3), while no such features were resolved on bare triangles. Moreover, nanostructures were incubated with TNKS-ARC4 (1:10 molar ratio, RT, 1 h) and analysed by mass photometry (Figure 5B-E). A positive mass shift was observed when TNKS-ARC4 was incubated with triangle-TBP, whereas no significant shift was observed for bare nanostructures incubated with TNKS-ARC4 (Figure 5C). Similarly, tetrahedron carrying TBP showed a significant shift when incubated with TNKS-ARC4, which was not observed for the bare tetrahedron (Figure 5D-E).

**Figure 5:**
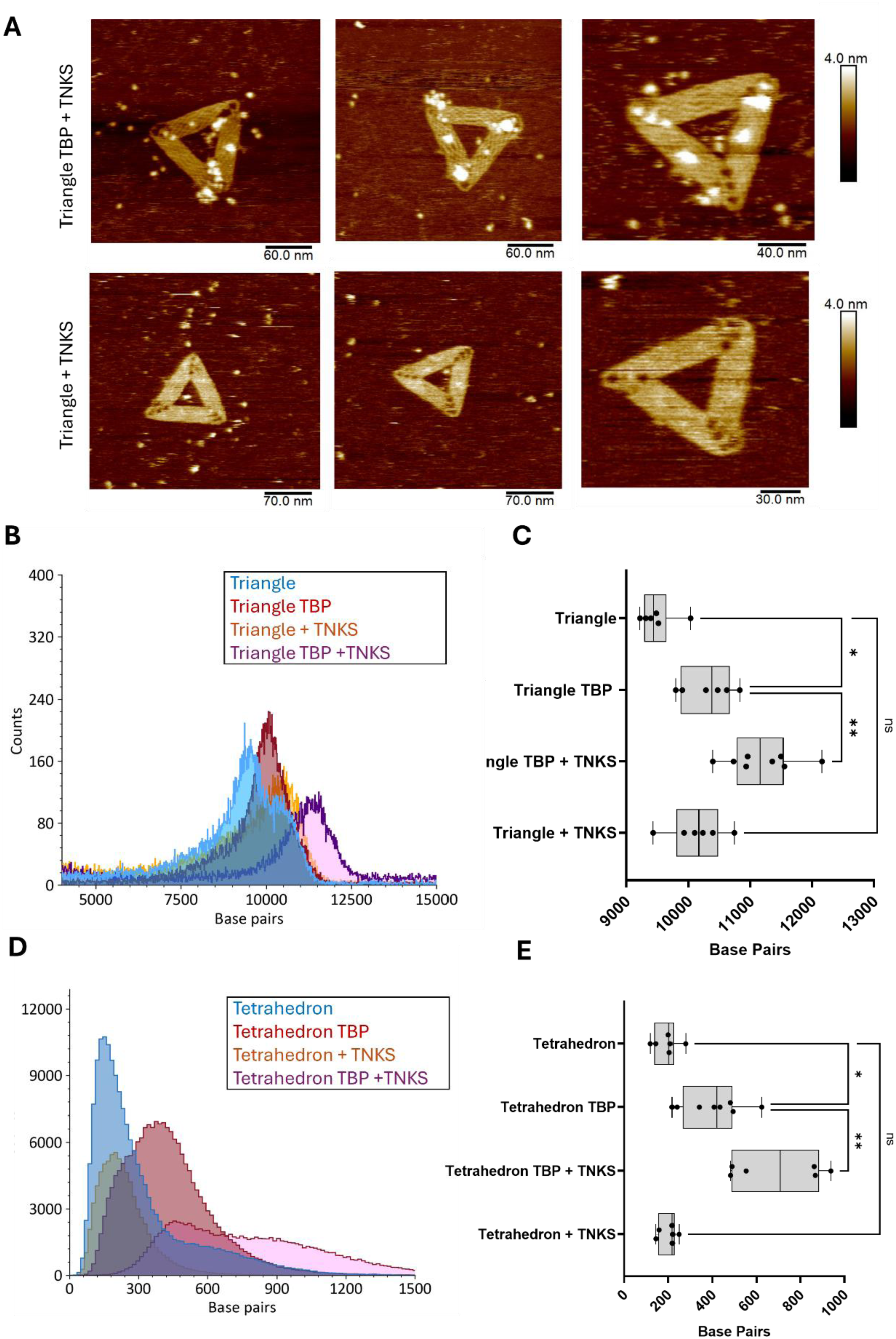
DNA nanostructures bind TNKS fragments in vitro. **A.** AFM images of triangles-TBP incubated with TNKS fragment show protein on the triangle surface. Protein is not detected on bare triangles **B**. Mass Photometry of triangle-TBP nanostructure in the presence of TNKS-ARC4 fragment shows TNKS binding. **C.** An average of three biological repeats of mass photometry shows significant shifts upon TNKS addition, corresponding to 18-21 molecules per nanostructure. **D.** Mass Photometry of tetrahedron-TBP nanostructure in the presence of TNKS-ARC4 fragment shows TNKS binding. **E.** An average of three biological repeats of mass photometry shows significant shifts upon TNKS addition to tetrahedron TBP.

### Nanostructure endocytosis via confocal microscopy

To assess cellular uptake of the DNA nanostructures, we first used HeLa Kyoto cells as a model cell line. HeLa cells possess a functional Wnt/β-catenin pathway that, unlike in colorectal cancer lines such as SW480, is not constitutively active: HeLa cells are wild-type for APC and β-catenin, so canonical Wnt signalling is basal but can be stimulated exogenously with Wnt3a. Cells were seeded in ibidi 8-well plates, treated with nanostructures (bare triangle, triangle-TBP, bare tetrahedron, tetrahedron-TBP, and free TBP) in serum-free OptiMEM, and imaged live at 3 h and 24 h post-incubation. Nanostructure concentrations were matched to equivalent TBP concentrations across conditions to ensure comparability (3 nM triangle nanostructures, equivalent to ∼100 nM TBP; 50 nM tetrahedron nanostructures, also equivalent to ∼100 nM TBP). Cells were stained with calcein-AM to visualise the cytosol and LysoBrite Orange to label lysosomes, while nanostructures were imaged in the 647 channel (AF647, Cy5).

Image analysis was performed using an in-house CellProfiler pipeline (Supplementary Figure S4). Briefly, cells were segmented based on the shape of cytoplasmic intensity, and debris was excluded by size filtering. The AF647 channel was smoothed using a bilateral filter, which limits Gaussian smoothing across edges to retain spatial resolution [46], then thresholded using the Robust Background algorithm (the dimmest 90% of intensity values were trimmed, and the threshold was set to 4 standard deviations above the mean of the remaining signal), with the lower threshold bound fixed at 0.08 to prevent spuriously low thresholds in buffer-only controls. Nanostructure puncta (origami clusters of varying size) were then segmented into primary objects based on local intensity differences between neighbouring clusters. The origami-buffer control returned zero detected puncta at any time point, confirming that the pipeline reliably discriminated nanostructure signal from background.

Triangle nanostructures exhibited moderate uptake at 3 h, with most puncta localised near the cell periphery (Figures 6, 8A) for both bare and TBP-functionalised triangles, with the latter showing consistently higher uptake; functionalisation with TBP therefore did not hinder, but rather enhanced, nanostructure endocytosis. Tetrahedral nanostructures were endocytosed more efficiently, particularly when TBP-functionalised (Figures 7, 8A), with signal again increasing significantly between 3 h and 24 h. Z-stack imaging confirmed that nanostructures were internalised rather than clustered on the cell surface (Supplementary Figure S7).

**Figure 6:**
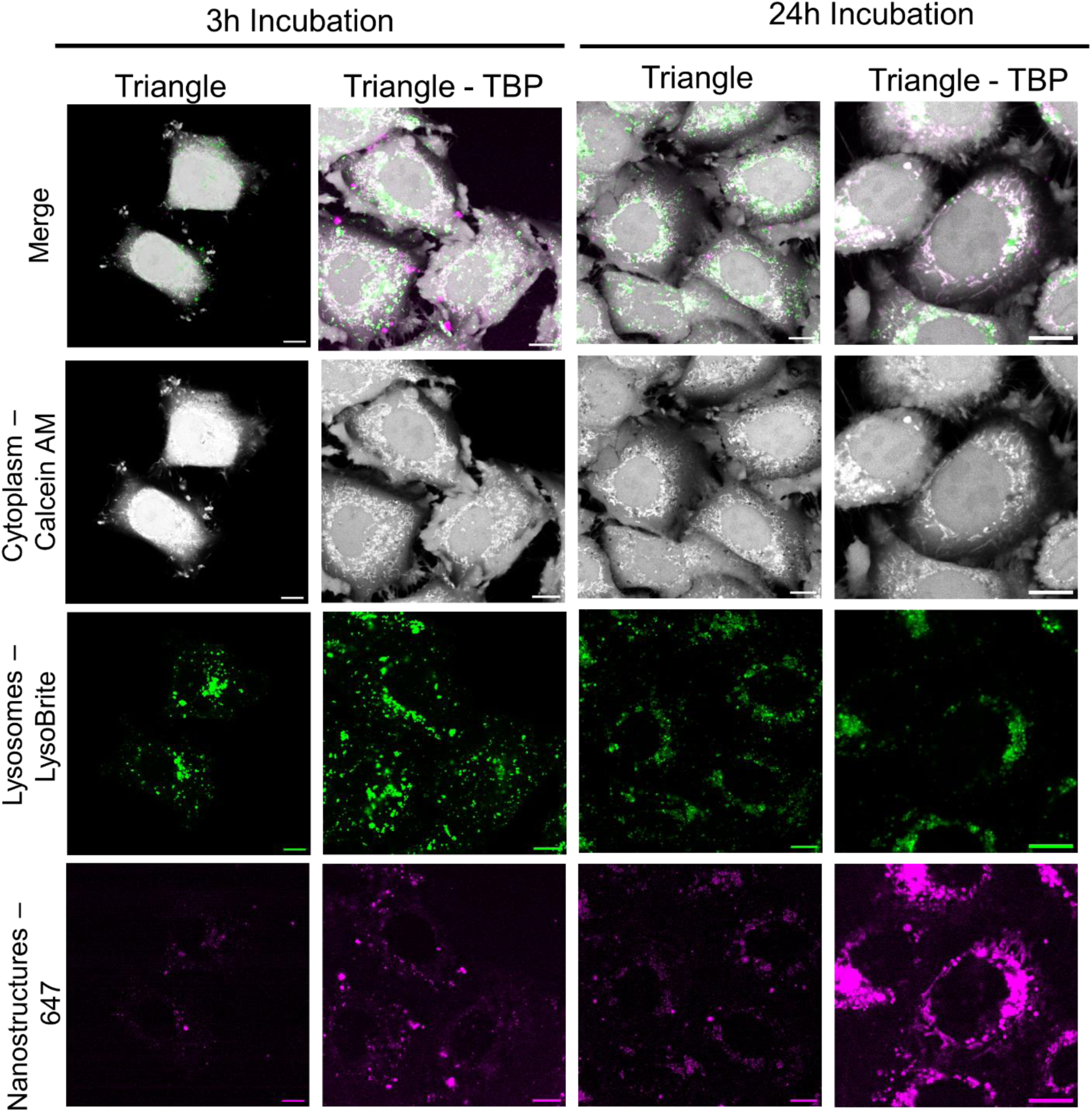
Selected confocal images of HeLa Kyoto cells incubated with bare and modified triangles, for 3h and 24h. Cells were stained with Calcein-AM and Lysobrite orange prior to imaging, nanostructures are detected in the 647 channel via AF647 on the nanostructures and Cy5 on the TBP. Scale bar corresponds to 10 μm. Larger frame images can be found in the supplementary (Figure S5).

**Figure 7:**
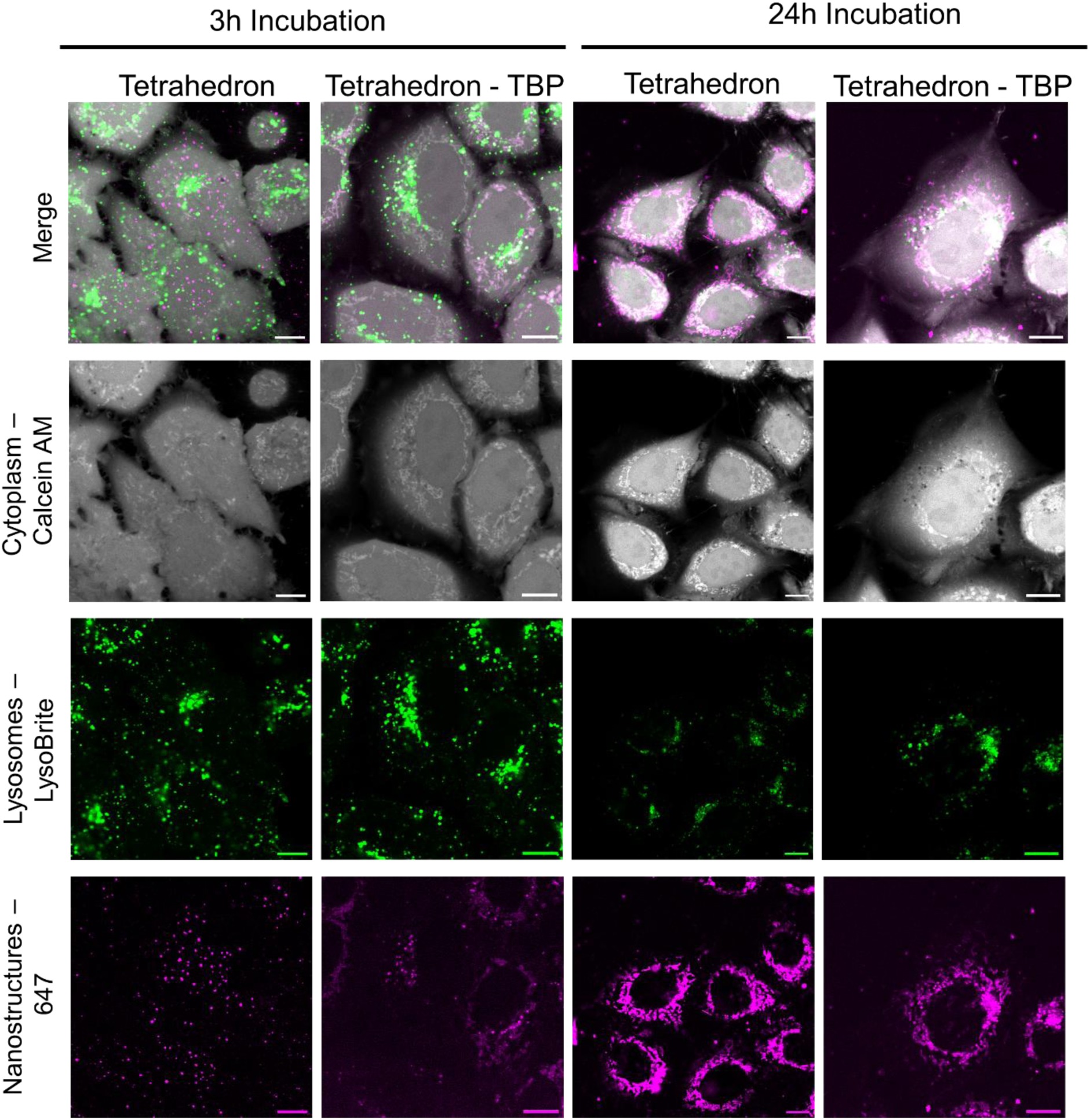
Selected confocal images of HeLa Kyoto cells incubated with bare and modified tetrahedra, for 3h and 24h. Cells were stained with Calcein-AM and Lysobrite orange prior to imaging, nanostructures are detected in the 647 channel via AF647 on the nanostructures and Cy5 on the TBP. Scale bar corresponds to 10 μm. Larger frame images can be found in the supplementary (Figure S6).

**Figure 8:**
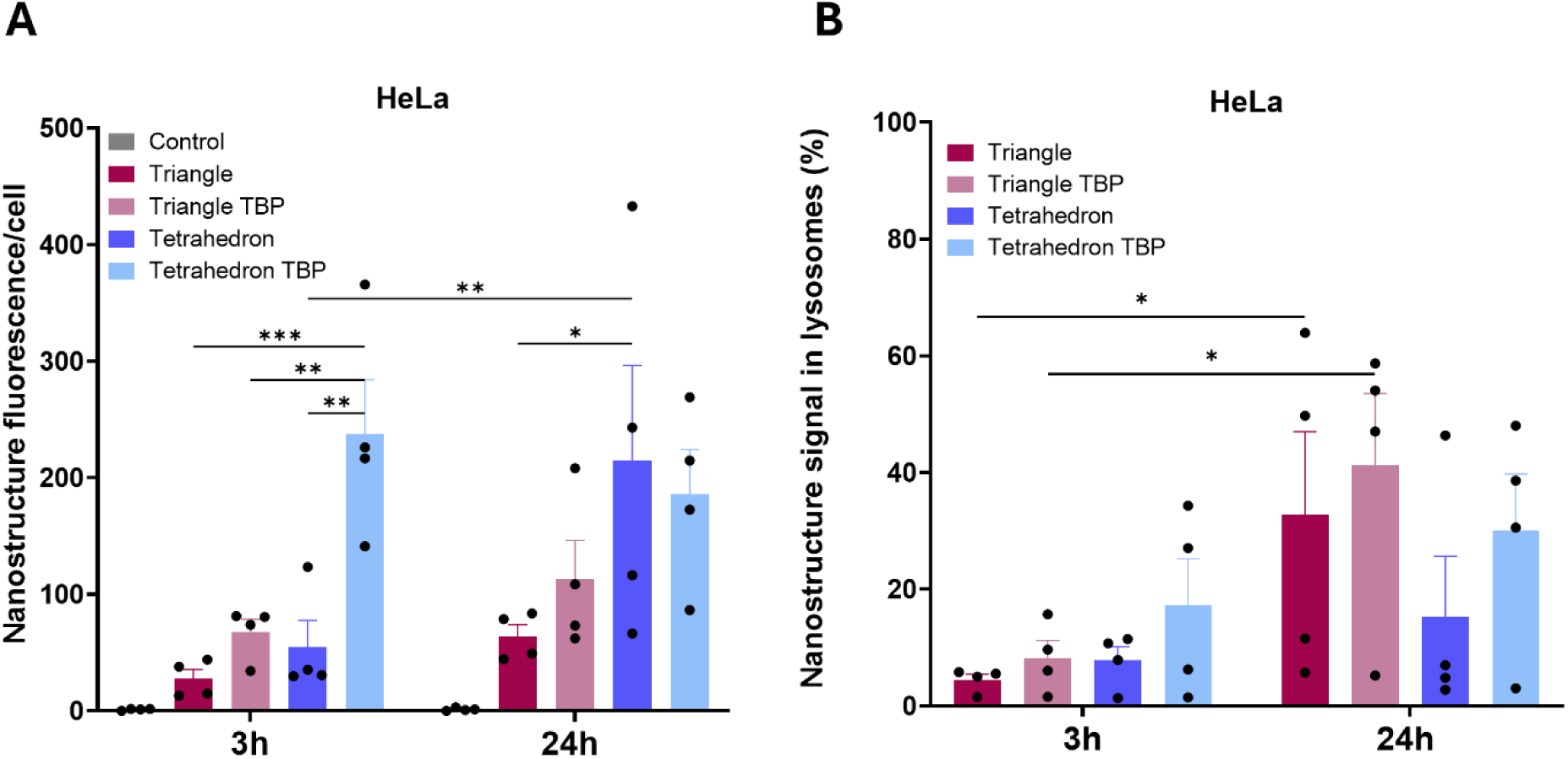
Image analysis quantification results for the HeLa Kyoto cell line. Conditions tested are triangle (bare, TBP-modified), tetrahedron (bare, TBP-modified), control (no treatment) and incubation times were 3 h and 24h. A. Nanostructure internalisation represented as number of nanostructure fluorescence puncta per cell, four biological repeats, n>50 cells per biological repeat. B. Percentage of lysosomal colocalisation. Mean of technical replicates is plotted for the percentage of Alexa647 signal in lysosomes relative to the whole cell. Bars represent standard error of mean for biological replicates, and a two-way repeated measures ANOVA was used to determine significance (p<0.05=*, p<0.01=**, p<0.001=***, p<0.0001=****). Analysis performed via in-house developed CellProfiler pipeline.

Lysosomal colocalisation was quantified as the integrated far-red signal spatially overlapping with LysoBrite signal, expressed as a percentage of total integrated signal per cell. All nanostructure treatments showed <60% colocalisation, indicating that a substantial proportion of internalised nanostructures remained in early endocytic compartments or potentially reached the cytosol. Free TBP was also internalised but exhibited markedly higher lysosomal colocalisation (Supplementary Figures S8, S9), consistent with rapid trafficking to degradative compartments in the absence of a protective scaffold.

We next extended the endocytosis study to SW480 colorectal cancer cells that stably expresses cytoplasmic GFP. SW480 cells express a truncated APC that leads to Wnt pathway dysregulation. Cells were seeded and treated identically to the HeLa experiments, with GFP signal used to visualise the cytosol and LysoBrite Orange used to label lysosomes, and nanostructures were detected again in the far-red channel.

Triangle endocytosis at 3 h was moderate, lower than in HeLa cells, and puncta predominantly localised at the cell periphery (Figure 9); in several cases, nanostructures were observed clustering along the outer cell membrane, outlining the cell boundary. By 24 h, internalisation increased significantly, both for bare and TBP-functionalised triangles, though overall signal remained lower than in HeLa cells. Tetrahedra showed a similar pattern to triangles, with moderate endocytosis at 3 h increasing further by 24 h, and membrane clustering again observed, particularly at the earlier time point. Lysosomal colocalisation was low across all conditions (<20%), suggesting that nanostructures may be predominantly cytosolic or in early endocytic compartments. As in HeLa cells, a fraction of free TBP was internalised but showed significantly higher lysosomal colocalisation than the nanostructures (Supplementary Figures S8, S9). Importantly, TBP functionalisation did not hinder uptake in any condition.

**Figure 9:**
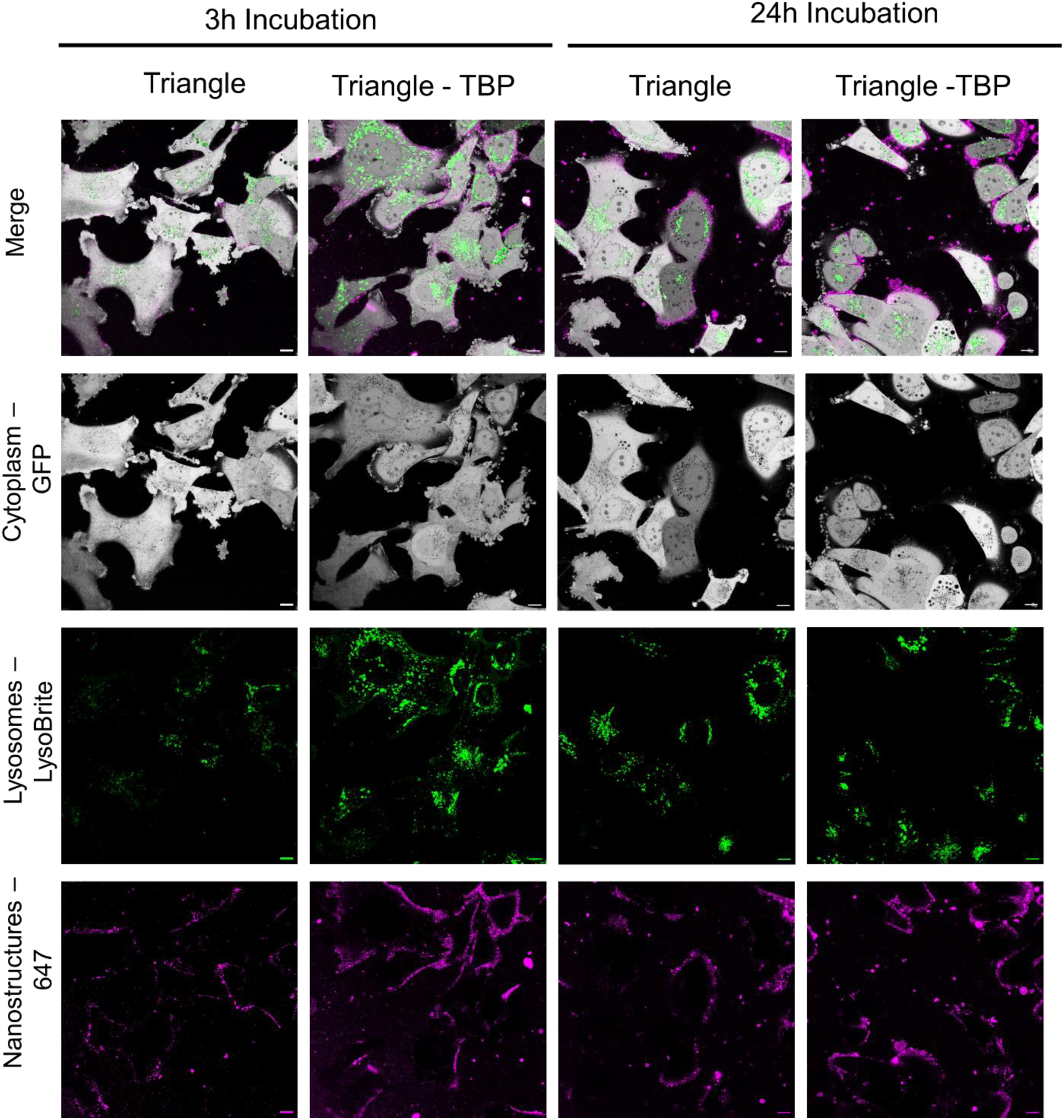
Selected confocal images of SW480 cells incubated with bare and modified triangle, for 3h and 24h. Stable cell line expresses cytosolic GFP. Cells were stained with Lysobrite orange prior to imaging, nanostructures are detected in the 647 channel via AF647 on the nanostructures and Cy5 on the TBP. Scale bar corresponds to 10 μm.

**Figure 10:**
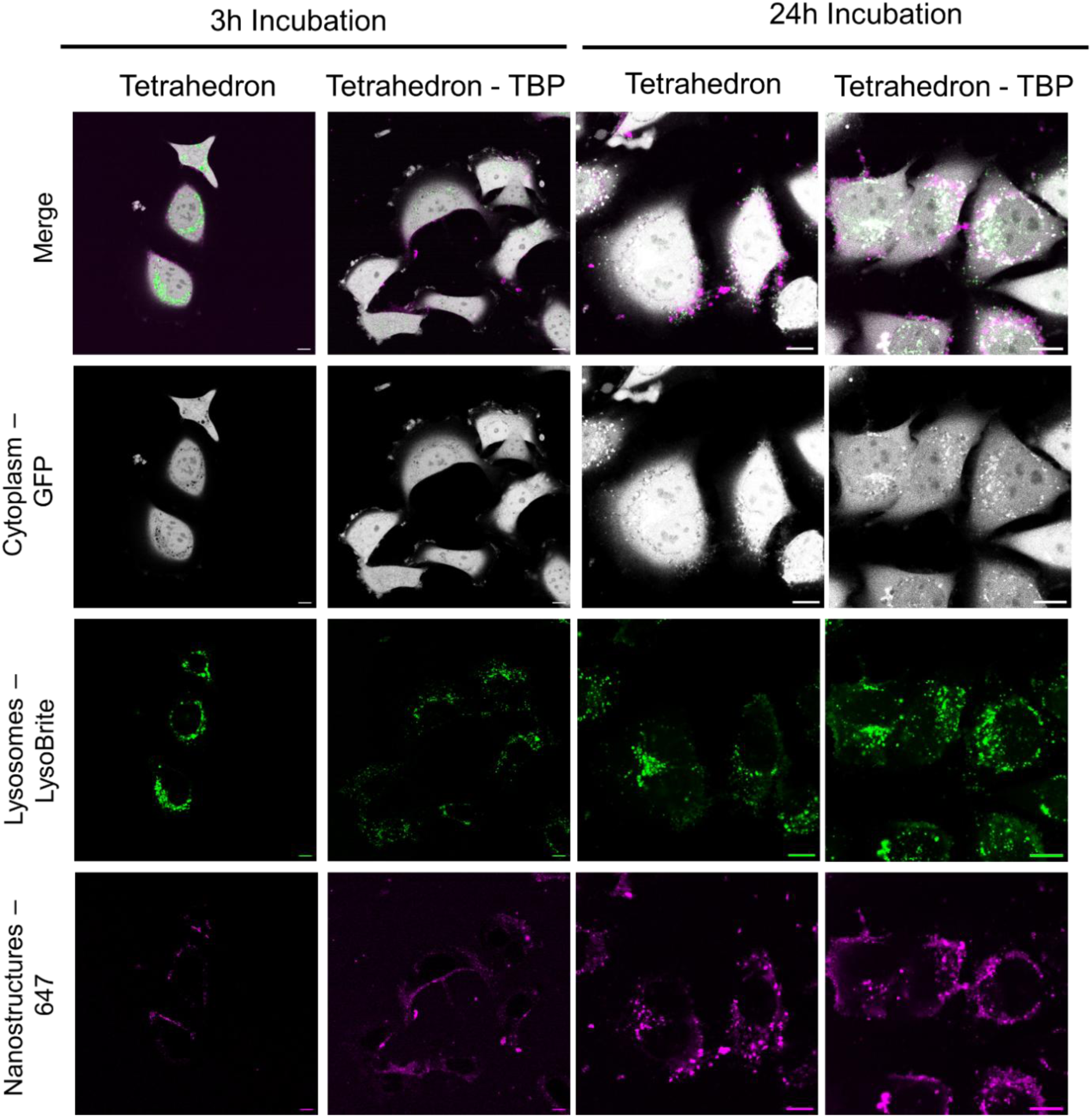
Selected confocal images of SW480 cells incubated with bare and modified tetrahedra, for 3h and 24h. Stable cell line expresses cytosolic GFP. Cells were stained with Lysobrite orange prior to imaging, nanostructures are detected in the 647 channel via AF647 on the nanostructures and Cy5 on the TBP. Scale bar corresponds to 10 μm.

**Figure 11:**
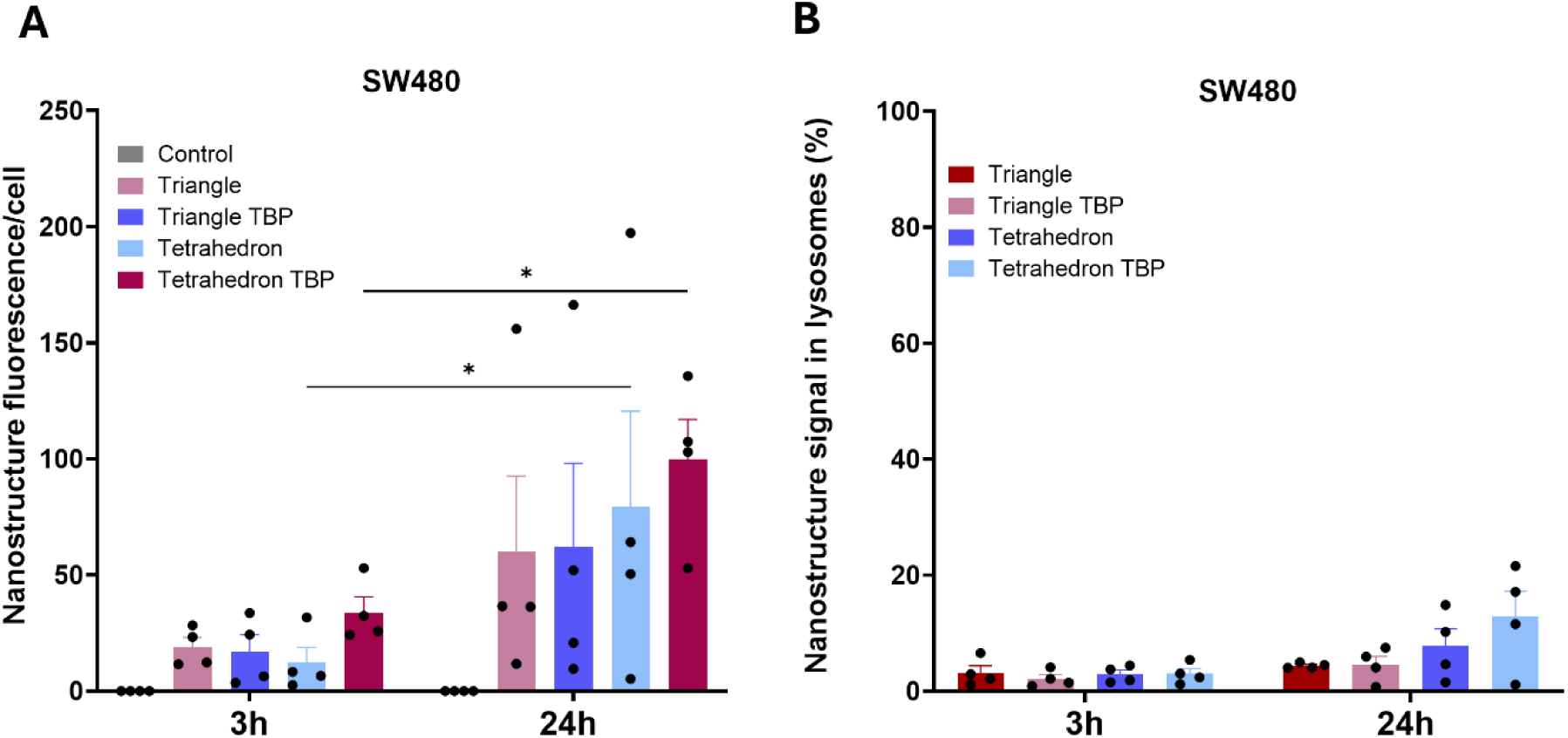
Image analysis quantification results for the SW480 cell line. Conditions tested are triangle (bare, TBP-modified), tetrahedron (bare, TBP-modified), control (no treatment) and incubation times were 3 h and 24h. A. Nanostructure internalisation represented as number of nanostructure fluorescence puncta per cell, four biological repeats, n>50 cells per biological repeat. B. Percentage of lysosomal colocalisation. Mean of technical replicates is plotted for the percentage of Alexa647 signal in lysosomes relative to the whole cell. Bars represent standard error of mean for biological replicates, and a two-way repeated measures ANOVA was used to determine significance (p<0.05=*, p<0.01=**, p<0.001=***, p<0.0001=****). Analysis performed via in-house developed CellProfiler pipeline.

### Functional effects on Wnt pathway activation

To determine whether TBP-functionalised nanostructures produce a functional effect on tankyrase following internalisation, we performed a Wnt luminescence reporter assay (TOPFlash). HeLa cells were transfected with a nanoluciferase reporter under a β-catenin-dependent, TCF/LEF-controlled promoter, whereas the reporter was integrated into SW480 cells. Cells were treated with nanostructures, free peptide, or small-molecule tankyrase inhibitors (IWR-1 and G007-LK) as positive controls. Nanostructure concentrations were again normalised to deliver equivalent effective TBP concentrations across conditions (3 nM triangle ≈ 100 nM TBP; 50 nM tetrahedron ≈ 100 nM TBP; 100 nM free TBP; 1 μM IWR-1; 1 μM G007-LK). For HeLa Kyoto cells, Wnt3a-containing media was added following nanostructure/inhibitor treatment. Viability was assessed using the cell-permeable fluorogenic substrate glycyl-phenylalanyl-aminofluorocoumarin (GF-AFC) to normalise for cell number, after which cells were lysed and nanoluciferase/firefly luciferase substrates added for luminescence measurement.

In HeLa Kyoto cells, treatment with triangle-TBP and tetrahedron-TBP reduced luminescence, indicating Wnt pathway downregulation, while bare nanostructures had no significant effect (Figure 12, Supplementary Table S1). The extent of downregulation was similar between triangle-TBP and tetrahedron-TBP. Free TBP at an equivalent concentration (100 nM) did not inhibit Wnt signalling. Small-molecule inhibitors also inhibited Wnt signalling, and to a greater extent than the nanostructure-TBP conditions; however, these were used at 1 µM, as recommended in the literature — tenfold higher than the effective TBP concentration in the nanostructure conditions. We have previously observed (Supplementary Figure S10) that lower concentrations of IWR-1 and G007-LK yield more moderate inhibition comparable to the nanostructure-TBP conditions.

**Figure 12:**
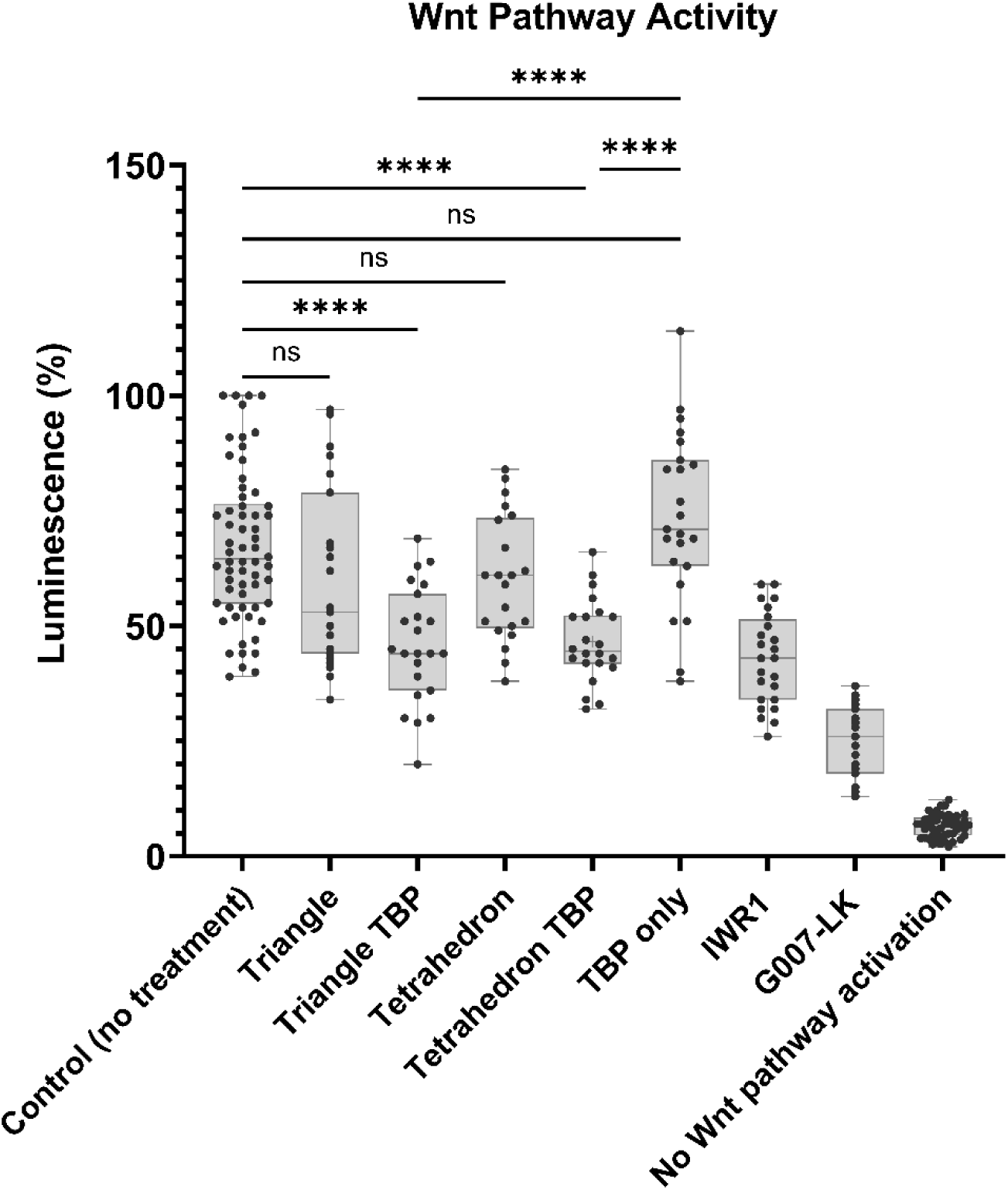
Wnt Assay results for HeLa Kyoto cell line after treatment with the nanostructures and small molecule TNKS inhibitors G007-LK and IWR1. Treatment concentrations are as follows: 3 nM triangle, triangle TBP (∼100 nM effective TBP concentration), 50 nM tetrahedron, tetrahedron TBP (∼100 nM effective TBP concentration), 100 nM free TBP, 1μΜ IWR1, 1 μM G007-LK, incubation time 24h in serum-free media. Data are average of four biological repeats (>24 technical repeats). Statistical significance was determined through a one-way ANOVA (p<0.05=*, p<0.01=**, p<0.0001=****). All comparisons can be found in Supplementary table S3.

In contrast, the Wnt assay in SW480 cells showed no significant differences across any treatment, including the small-molecule inhibitors (Supplementary Figure S11, Supplementary Table S2). This may be partially attributable to reduced nanostructure internalisation in this cell line relative to HeLa; however, the lack of effect from small-molecule inhibitors as well points to an additional mechanistic explanation. SW480 cells carry a truncating APC mutation that removes the AXIN-binding repeats required for a fully functional β-catenin destruction complex, so it is unsurprising that TNKS inhibition alone is insufficient to suppress Wnt signalling in this context.

## Discussion

In this study, we demonstrate that DNA nanostructures functionalised with a tankyrase-binding peptide (TBP) can be reproducibly assembled, retain the ability to bind TNKS *in vitro*, are internalised by mammalian cells, and in HeLa Kyoto cells produce measurable downregulation of Wnt/β-catenin signalling. By presenting a tankyrase-binding peptide (TBP) on two distinct DNA architectures—a high-valency 2D triangle and a compact, low-valency 3D tetrahedron—we show that both geometries can be efficiently functionalised, internalised by mammalian cells, and engage TNKS to produce a measurable biological effect.

The tetrahedron’s superior endocytosis relative to the triangle in both cell lines is in line with the wider nanoparticle literature, in which uptake efficiency via receptor-mediated endocytosis declines toward the upper size limit for clathrin-mediated entry (∼200 nm) [48], [49]. The ∼10 nm tetrahedron falls within a favourable size range, while the ∼100 nm triangle is expected to be less efficient. Shape is another factor: DNA-origami-specific studies comparing nanostructures of similar mass have found that compact, caged 3D geometries are internalised more efficiently than flat, sheet-like, or high aspect-ratio structures [25], [33], [34] and the triangle’s large, low-aspect-ratio 2D geometry may present a fundamentally different interface to scavenger receptors and the plasma membrane than a compact 3D tetrahedron, independent of its overall mass [32]. Together, size and geometry likely both contribute to the more efficient uptake of the tetrahedron observed here. Another notable observation was that TBP functionalisation consistently enhanced, rather than hindered, endocytosis of both geometries. This raises the possibility that the peptide itself, its Cy5 label, or the added surface charge/hydrophobicity of the conjugate favourably alters interactions with the cell surface. The membrane-associated clustering we observed for both nanostructures in SW480 cells at early time points, but not prominently in HeLa cells, may reflect differences in scavenger receptor expression or surface composition between the two lines; this remains speculative and would benefit from direct receptor-expression profiling.

Functional Wnt pathway downregulation by triangle-TBP and tetrahedron-TBP in HeLa cells is indirect evidence that a fraction of the internalised nanostructures reaches cytosolic TNKS. TNKS is not accessible from within the endo-lysosomal lumen, so pathway inhibition requires that TBP-bearing nanostructures, or a released TBP fraction, gain access to the cytoplasm. This is a notable result, since achieving a functional cytosolic readout from a nanostructure carrying no dedicated endosomal-escape-promoting moiety (e.g. pH-responsive elements, or lipidated anchors) remains a widely recognised bottleneck [50], [51]. Both nanostructures showed substantial non-lysosomal signal (<60% colocalisation in HeLa; <20% in SW480), which we interpret as consistent with a meaningful cytosolic and/or early-endosomal fraction.

Despite the tetrahedron being internalised to a greater extent than the triangle in HeLa cells, triangle-TBP and tetrahedron-TBP produced a similar degree of Wnt pathway downregulation. Taken together, these two observations suggest that, on a per-nanostructure basis, the triangle achieves a comparable functional effect from substantially less internalised material. This implies that once internalised, the triangle is the more potent inhibitor of the two geometries. This fits with its higher TBP valency: presenting 27 copies of TBP, rather than 2, is expected to confer a proportionally greater avidity advantage for engaging TNKS, which is itself a multivalent, clustering target. This mechanistic interpretation is consistent with the broader avidity literature, in which binding advantage does not scale linearly with ligand copy number but can continue to increase with valency beyond a low threshold, particularly against clustered or multivalent targets [52], [53], [54]. It must be noted, however, that this comparison rests on combining two independently measured, semi-quantitative readouts (fluorescence puncta counts and luminescence signal) and should be treated as an interpretation of the pattern in our data rather than a precisely quantified potency difference.

Free TBP at an equivalent concentration (100 nM) had no effect on Wnt signalling in either experiment. We attribute this observation to two factors: as in our previous work using CTPR-scaffolded multivalent TBP constructs [16], very high concentrations of free peptide were needed to see an effect on the Wnt pathway and compete with the endogenous substrate pool - functional inhibition appears to require multivalent presentation, not TBP delivery alone. Second, free TBP showed substantially higher lysosomal colocalisation than either nanostructure suggesting that a smaller fraction of the internalised peptide is available in the cytosol to begin with. The absence of a functional effect from free TBP is therefore likely a consequence of both insufficient avidity and reduced cytosolic accessibility, rather than either factor alone.

The absence of any Wnt-inhibitory effect of either small-molecules or DNA nanostructures in SW480 cells likely points to a genotype effect on TNKS-inhibitor efficacy. Sensitivity of colorectal cancer cells to TNKS inhibition depends on the position of the truncating APC mutation relative to the SAMP AXIN-binding repeats. Short-form truncations that remove all repeats, as in SW480, fail to support efficient activity of the destruction complex even when AXIN is stabilised [55], [56], [57]. AXIN stabilisation is therefore insufficient to impact destruction complex activity in SW480 cells. Moreover, our experiments in the two cell lines differ in another way: in HeLa cells, nanostructures and inhibitors were added before Wnt3a stimulation, so the functional effect measured is prevention of pathway activation, not reversal of an already-active pathway, comparing between a prophylactic outcome and an interventional one.

Overall, we observed here that multivalent presentation of TBP, rather than the free peptide itself, was required to achieve effective TNKS inhibition and Wnt pathway downregulation in HeLa cells. This held across two geometrically distinct nanostructures presenting an order-of-magnitude difference in TBP copy number, indicating that the underlying design principle is not restricted to a single scaffold architecture. The fact that this functional effect was achieved without any dedicated endosomal-escape-promoting modification indicates that a sufficient fraction of the internalised nanostructures reaches the cytosol to engage TNKS. The ability to engineer multivalent inhibitors with defined nanoscale geometry opens new opportunities for modulating signalling pathways governed by clustering, polymerisation, or phase separation.

## Methods

### Generating peptide conjugates

TBP-linker conjugates were generated through catalyst-free Click reaction. Linker X (TTTTTCATAATAACTCTTAGATTTG-azide) (100 μl, 100 μM) (IDT) was added to TBP peptide (DBCO-(Peg)2-C(Cy5)NREAGDGEE) (100 μl, 100 μM) (Peptide Synthetics). Mixtures were incubated at 4℃ in rolling incubator overnight. Click reactions were validated with 20% urea acrylamide gels. Gel was electrophorized in 0.5x preheated TBE buffer for 5 minutes at 100 V and then 15 minutes for 300 V. The gel was stained with 1/50000 SYBR^TM^ Safe (Invitrogen) for 15 minutes prior to UV imaging with the Gel Doc^TM^ EZ imager (Bio-Rad). Conjugates were purified with HPLC using the Clarity 5 μm Oligo-RP column (Phenomenex) with 100 μl injections at a flow rate of 1.5 ml/minute.

### Synthesis and purification of DNA origami

M13mp18 DNA scaffold (10 μl, 100 nM) was added to staple mix (25 μl, 200 nM) in 10 μl 10x OB and 55 μl MilliQ water. The mixture was heated to 90℃ in a thermocycler (ThermoFisher) and then cooled at a rate of -1℃/minute to 25℃. To hybridise the nanostructures, 10x molar excess of conjugate was added to DNA nanostructures and hybridised by heating to 40℃ for 60 minutes followed by cooling at -1℃/minute to 15℃. DNA nanostructure formation was confirmed with a 1% agarose gel which was precast with 1x TBE supplemented with 5 mM MgCl_2_ Buffer and 5 μl SYBER^TM^ Safe (ThermoFisher). The gels were electrophorised on ice at 90 V for either 45, 90, or 190 minutes.

DNA nanostructures without conjugate functionalisation were purified from excess staples by size exclusion chromatography. Sephacryl S-300 (Cytiva) columns were packed according to manufacturer’s instructions and the DNA origami sample was eluted by centrifugation at 1000 g for 4 minutes [58]. Conjugate-hybridised DNA nanostructures were centrifuge purified at 20 minutes at 9,600 g, and the sample was concentrated by resuspending the pellet in 1x OB at 1/10 of the original sample volume.

### Atomic Force Microscopy

Purified DNA origami diluted 1:10 in 20 μl 1x OB were adhered to a cleaved mica. Imaging was performed in 1x OB on the Dimension FastScan AFM (Bruker) with FastScan-D probes. Tapping mode in fluid was used with 512 samples collected per line and a scan rate of 17.5 Hz. Images were processed with Nanoscope 9.7 software.

### Mass photometry

15 μl of 20 mM Mg^2+^ origami buffer was placed in a 6-well silicone gasket and buffer dilution calibration was performed on the Refeyn TwoMP. Then 5 μl of origami sample was loaded for measurement. The DNA nanostructure mass calibration was done according to manufacturer’s (Refeyn) instructions with a low mass DNA ladder (Invitrogen) in PBS. To assess TNKS binding, 2ul of 1mM TNKS stock was incubated with 5ul of 30 nM origami for 30 minutes. 13 μl of buffer was added, and a buffer calibration was performed prior to addition of the TNKS-origami mixture.

### Cell culture

SW480 active cells (Amsbio) were cultured in a T25 flask with DMEM supplemented with 10% FCS and 1% penicillin-streptomycin. The media was changed every 2 days and passaging occurred every 4 days. At ∼80% confluency, cells were washed once with PBS, incubated with trypsin-EDTA for 3 minutes, neutralised with supplemented DMEM, and split 1:4 into new T25 flasks. HeLa Kyoto cells were cultured and passaged as previously described except passaging occurred every 3 days. Cell cultures were not maintained for more than 20 passages.

### Confocal microscopy and image analysis

SW480 active cells were seeded into uncoated 8-well ibidi plates at a density of 3 × 10^4^ cells/well in DMEM supplemented with 10% FBS and 1% penicillin-streptomycin. After overnight attachment, wells were washed with 300 μl PBS and stained with Lysobrite Orange (1/2000) (AA5 Bioquest) in OptiMEM media and incubated for 20 minutes at 37℃ with 5% CO_2_. Media was aspirated and replaced with 180 μl OptiMEM mixed with 20 μl origami (30 nM), 1 μM TBP-Cy5, or 1x OB. Live cell imaging was performed on the STELLARIS Confocal Microscope (Leica) with the 63x oil objective at 37℃ and 5% CO_2_. Cells were imaged in the green channel (ex: 488 nm, em: 507 nm), red channel (543/565 nm), and far-red channel (649/667 nm) using frame sequential imaging, with a line average of 2 for each channel. 2D confocal images were processed using FIJI

Confocal images were analysed with an in-house CellProfiler (4.2.8) pipeline. Cells were segmented based on shape and GFP/Calcein intensity, and debris below a minimal area were excluded. The far-red channel was smoothed with a bilateral smoothing filter and the Robust Background algorithm was used to discern genuine from background signal. The dimmest 90% of intensity values were trimmed, and the threshold was set to 4 standard deviations above the mean of the remaining signal. Nanostructure peaks were distinguished by intensity differences and assigned to a cell if 95% of the signal spatially overlapped. To quantify lysosomal co-localisation, the total thresholded far-red signal within lysosomes was normalised to the total intracellular signal. Analyses of SW480 and HeLa cell images did not differ, except for cell segmentation parameters which were fine-tuned for each cell line.

### Wnt assays

Wnt assays were performed on firefly and nanoluciferase transfected HeLa Kyoto and SW480 cells. 10 μl of 30 nM origami and 10 μM IWR-1/G007-LK, and 1 μM TBP was added to each well. After another 8 or 32 hours 20 μl Wnt conditioned media obtained from L-cells expressing Wnt3A (ATCC CRL-2647) was added. After a 16-hour incubation, 80 μl of GF-AFC (1:500) in viability assay buffer was added to each well. After a 30-minute incubation, excitation/emission at 390/505 nm was determined with the CLARIOstar Plus microplate reader (BMG Labtech). Cells were lysed and prepared for luminescence measurement according to manufacturer’s instructions for the Dual Luciferase Kit (Promega).

### TNKS-ARC4 purification

His-tagged Tankyrase-2 Ankyrin Repeat Cluster 4 (residues 488-649, referred to as TNKS-ARC4) was expressed and purified as follows. The pRSET vector encoding the His-tagged construct was transformed into chemically competent Escherichia coli C41 cells. Colonies were grown at 37 °C in 2xYT media (Formedium) containing ampicillin (50 μg mL−1), shaking at 220 rpm until the optical density at 600 nm was ∼0.6. Protein expression was induced with 0.5 mM isopropyl β-d-1-thiogalactopyranoside (IPTG) for 16–20 hours at 20 °C. Cells were harvested and resuspended in buffer A (50 mM Tris–HCl, 150 mM NaCl, pH 8.0) including EDTA-free SIGMAFAST protease inhibitors (Sigma-Aldrich) and DNase I. Cells were lysed by high-pressure homogenization using an Emulsiflex C5 homogenizer (Avestin) at 15 000 psi, and cell debris was removed by centrifugation steps at 35 000g for 30 minutes. TNKS-ARC4 was purified by immobilised metal ion affinity chromatography (IMAC) on a 5 mL HisTrap Excel column according to the manufacturer’s instructions (Cytiva). The column was washed with 20 CV of buffer A containing 20 mM imidazole to prevent nonspecific interaction of lysate proteins to the beads. Proteins were eluted with buffer B (50 mM Tris–HCl, 150 mM NaCl, 500 mM imidazole, pH 8.0). Purified protein was flash-frozen and stored at −80 °C until further use. To remove the His tag, proteins were incubated with 125 U of thrombin (Sigma-Aldrich) overnight at RT on a rotating mixer.

## Acknowledgments

MZ acknowledges funding from the Ernest Oppenheimer Early Career Research Fellowship (School of the Physical Sciences, University of Cambridge).

IM acknowledges funding from the Royal Society (URF/R1/221795, IES∖R3∖223128, IES∖R2∖222107, RGS∖R1∖231266) and the National Biofilms Innovation Centre (BB/R012415/1 03PoC20-105).

LSI acknowledges funding into her lab from BBSRC (BB/Y007816/1) research grant.

SQ acknowledges support from the Robert Henderson Summer Internship programme (Emmanuel college, University of Cambridge).

We thank Dr Zutao Yu for assistance with HPLC measurements. Dr Pamela J. Rowling for useful discussions and advice on Wnt pathway and Biophysics, as well as TNKS-ARC4 purification. Dr Emmanuel Derivery for useful discussions on mechanisms of endocytosis and Cell Biology. Dr Katherine Stott for Mass Photometry advice and assistance, Dr Stephen McLaughlin for Biophysics advice, Dr Andrew Baker for nanoparticle discussions, and Dr Federico Bosetto for DNA nanotechnology discussions.

## Notes

### Competing Interest Statement

The authors have declared no competing interest.

### Summary of Updates

Author list and affiliations updated in the new version

