## Supplementary Information for "Programmable DNA-peptide nanostructures for multivalent regulation of intracellular signalling"

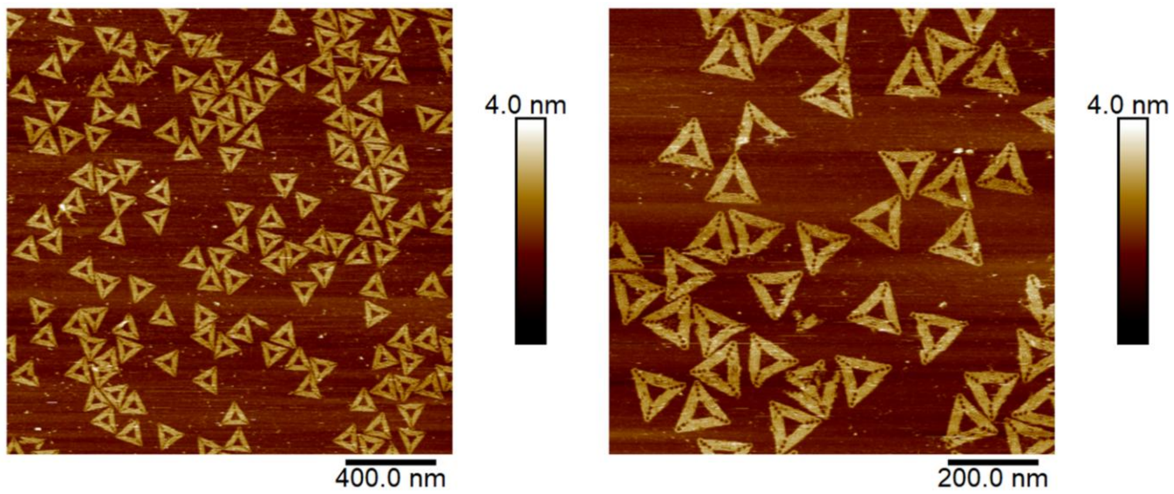

**Figure S1:** AFM images of unmodified (bare) triangle, showing distinct trapezoidal domains (~100 nm across the outer perimeter) connected to one another by bridging staples.

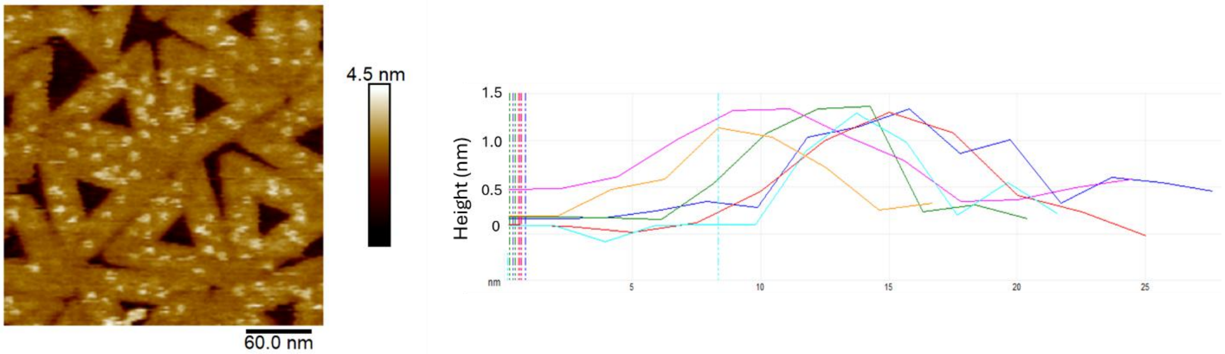

**Figure S2:** Cross-section of modification points on the triangle surface have an average height of ~1.5 nm. Six representative traces are shown.

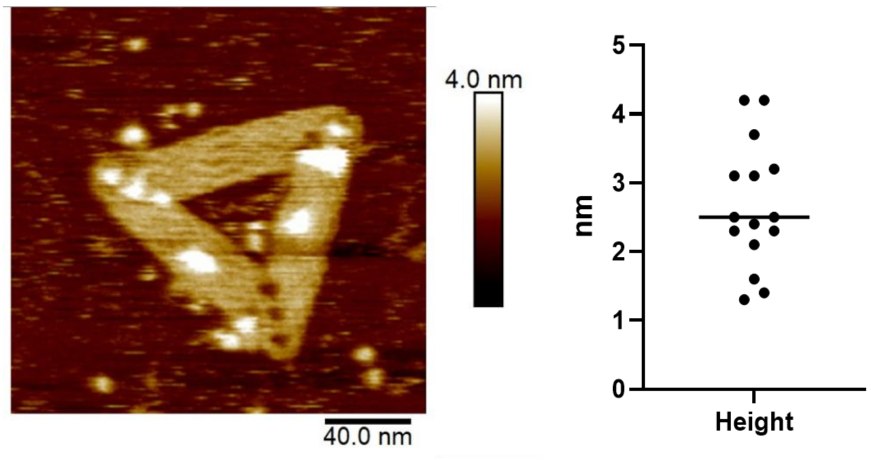

**Figure S3:** Average size of spherical features on the triangle surface after TNKS incubation is 3 nm, corresponding to a 25-30 kDa protein

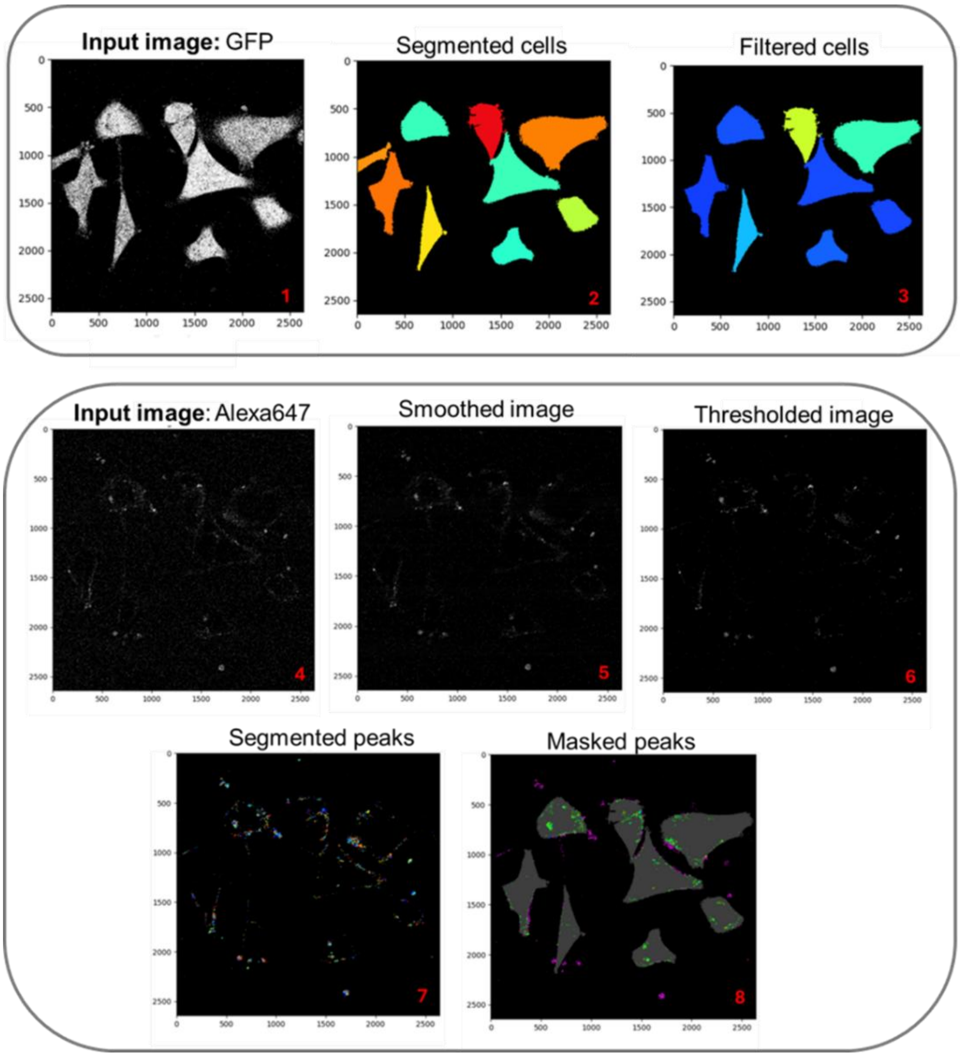

**Figure S4:** Schematic of in-house developed Cell Profiler pipeline for image analysis (cell segmentation and Alexa647/Cy5 peak identification).

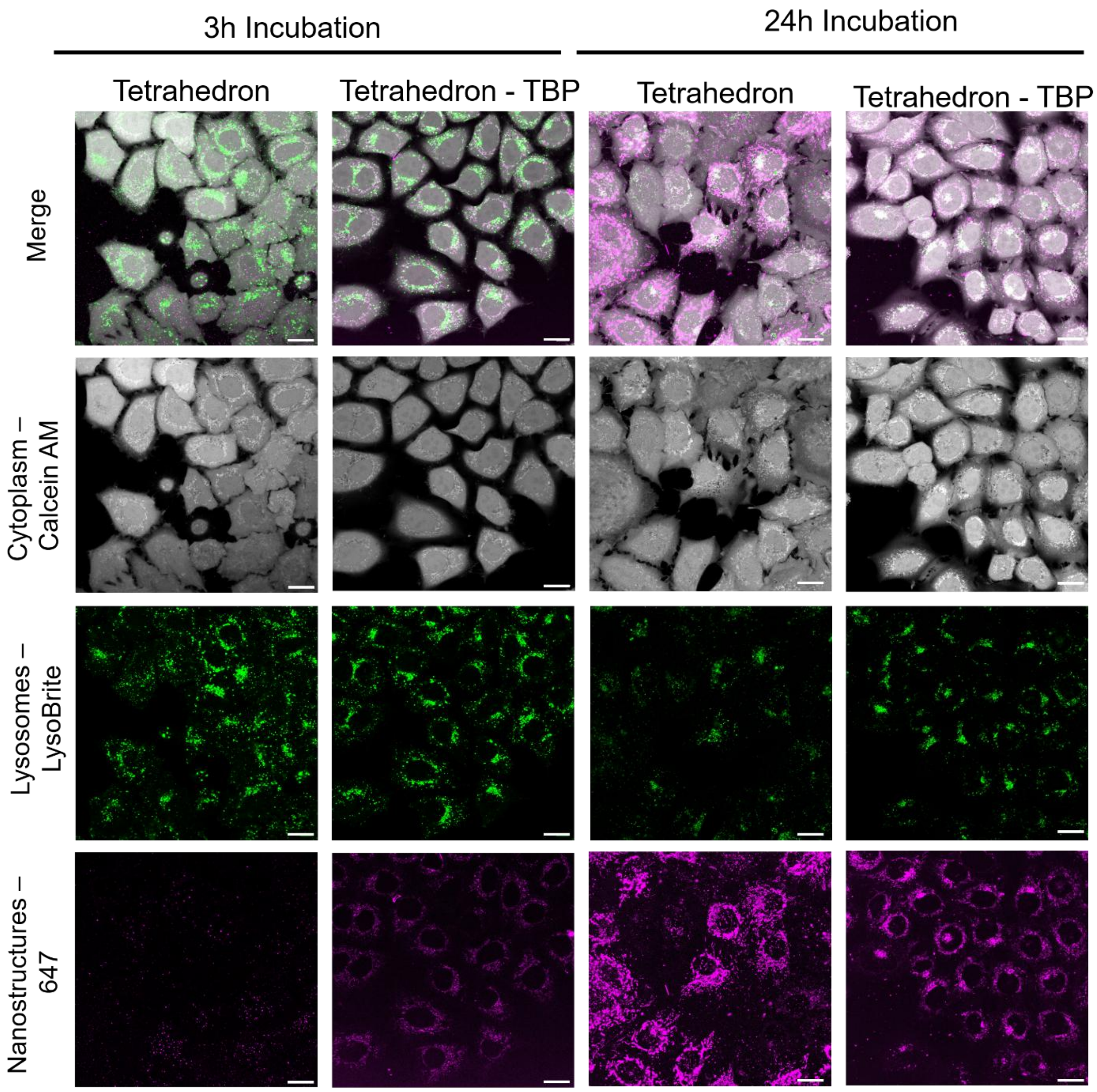

**Figure S5:** Endocytosis of tetrahedral nanostructures in HeLa Kyoto cells. Cells were stained with Calcein-AM and Lysobrite orange prior to imaging, nanostructures are detected in the 647 channel via AF647 on the nanostructures and Cy5 on the TBP. Scale bar corresponds to 20 μm.

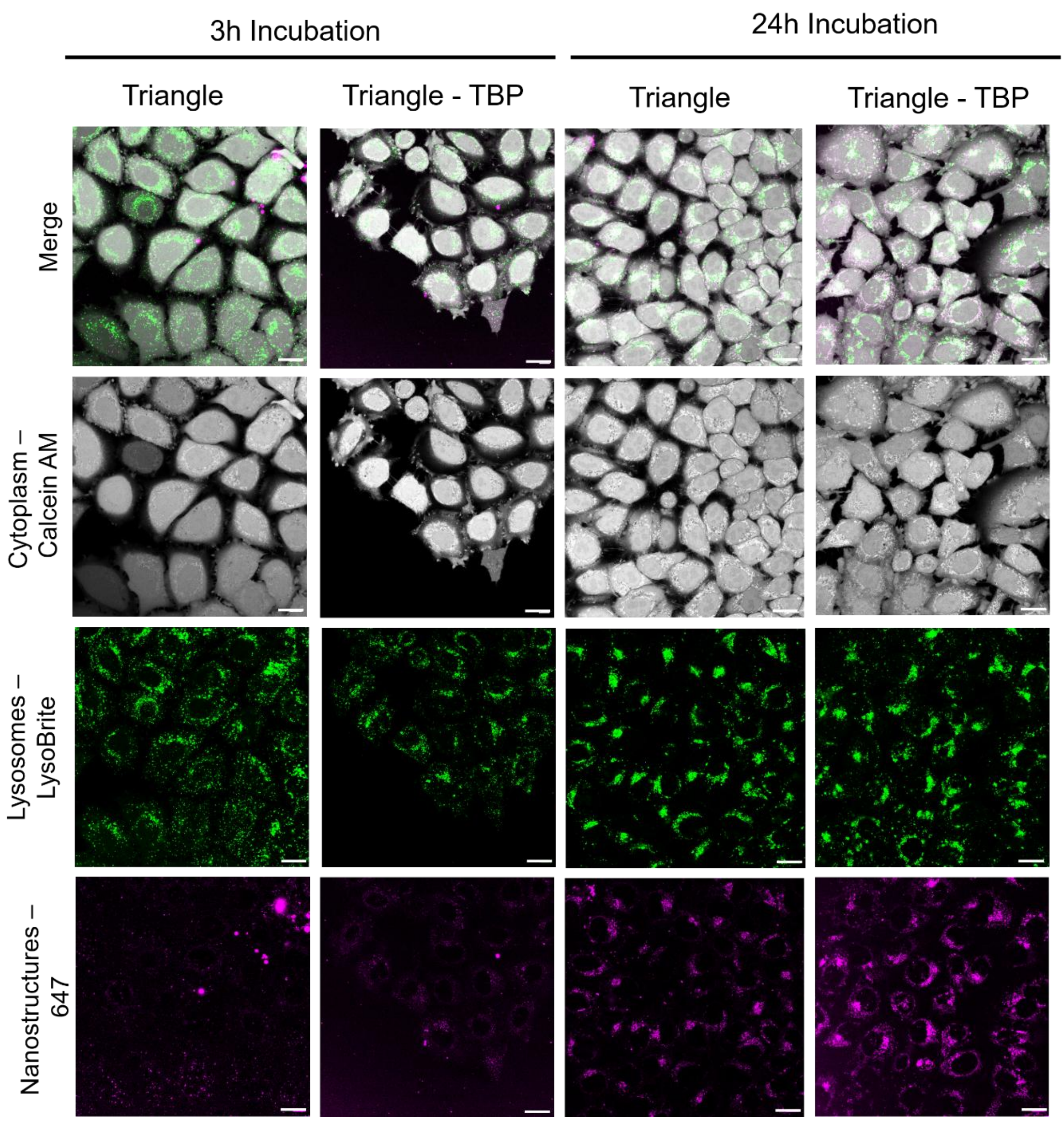

**Figure S6:** Endocytosis of triangle nanostructures in HeLa Kyoto cells. Cells were stained with Calcein-AM and Lysobrite orange prior to imaging, nanostructures are detected in the 647 channel via AF647 on the nanostructures and Cy5 on the TBP. Scale bar corresponds to 20 μm.

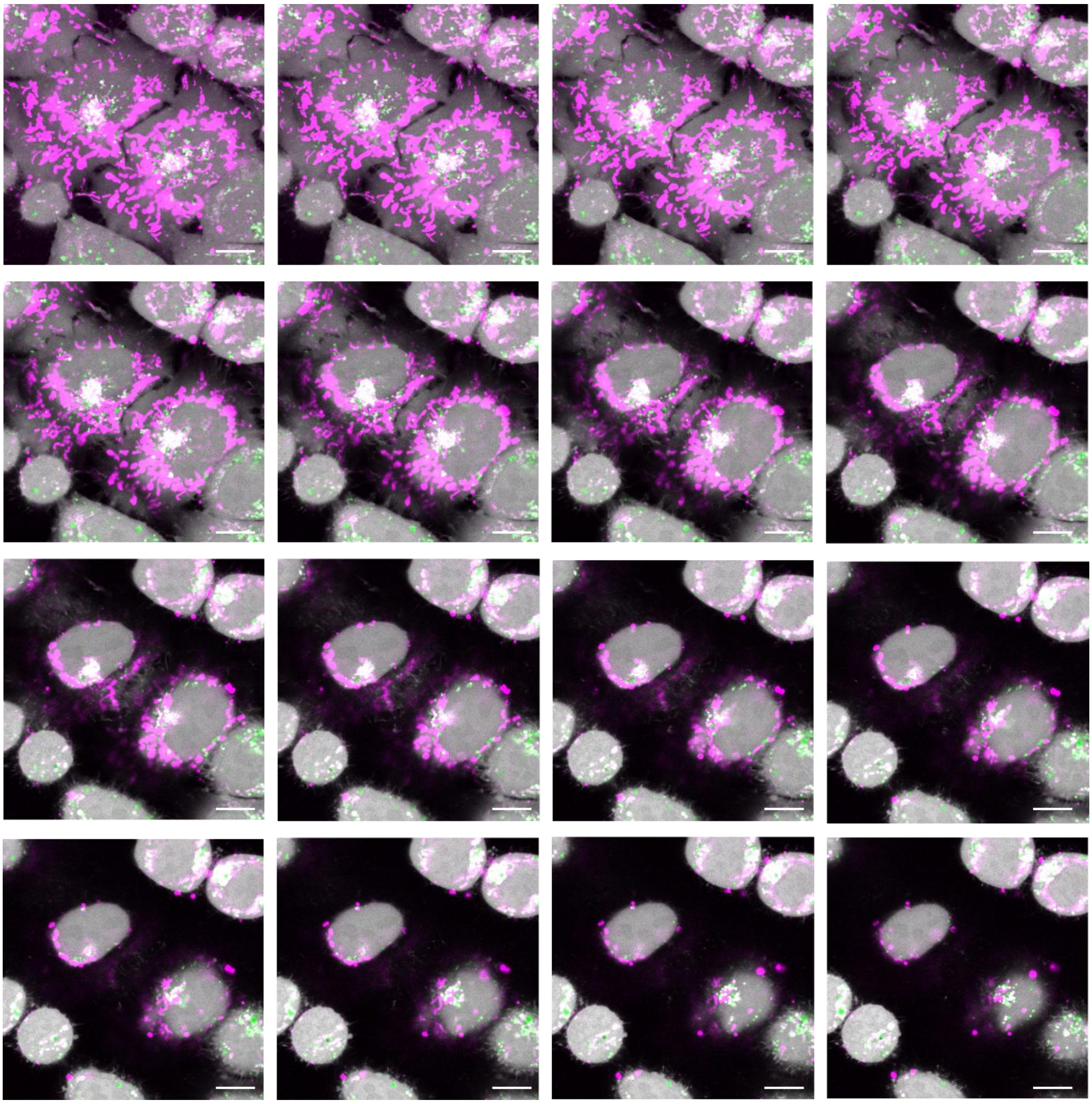

**Figure S7:** Representative Z-stack of HeLa Kyoto cells, 16 slices. Cells are treated with bare tetrahedron, 24h. Grey: cytoplasm, calcein AM, green: lysosomes, LysoBrite Orange, magenta: tetrahedron, AF647. The nanostructure signal can be detected inside the cells, rather than on their surface.

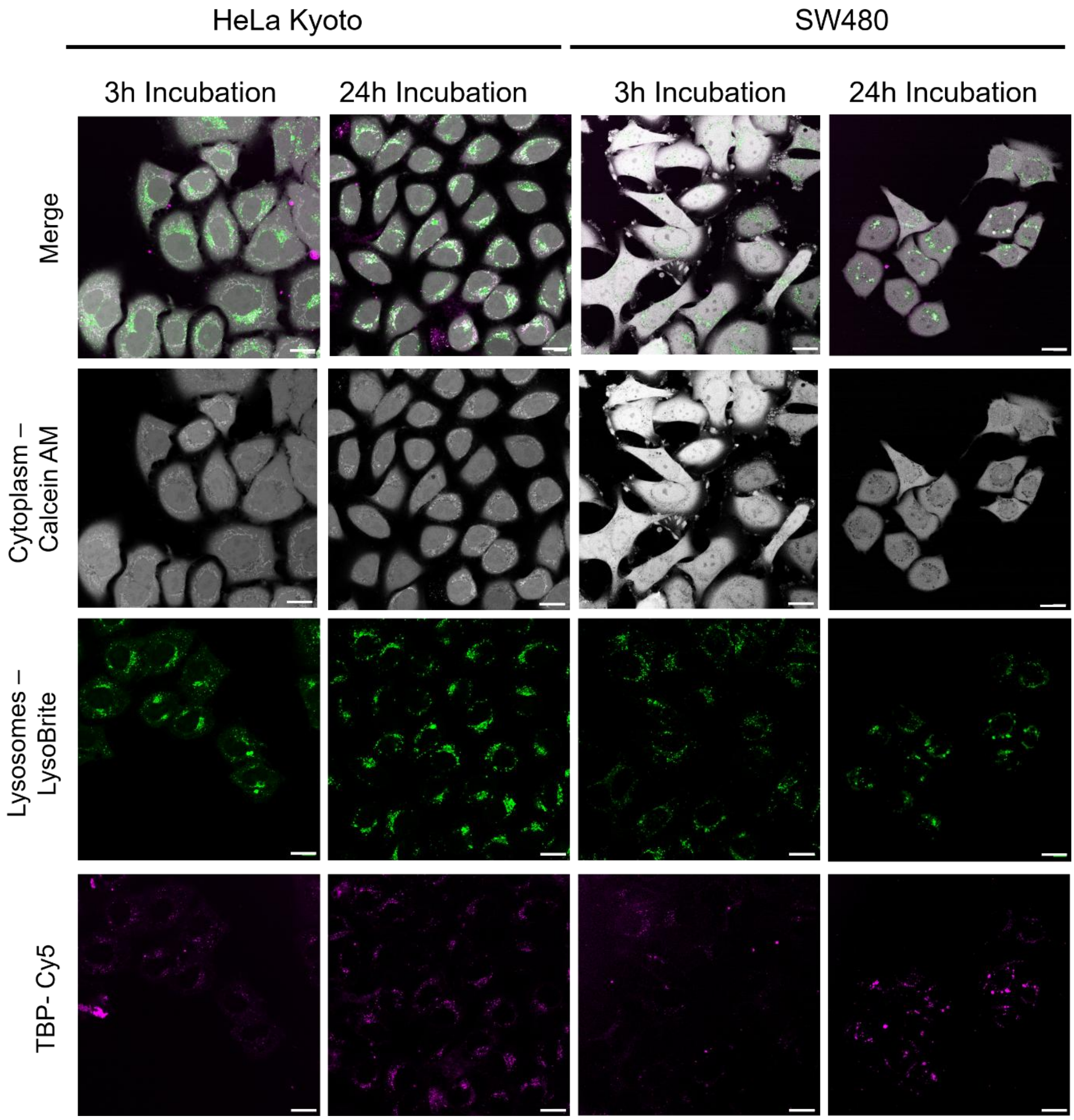

**Figure S8:** Endocytosis of TBP in HeLa Kyoto cells and SW480 cells. SW480 stable cell line expresses cytosolic GFP, HeLa cells were stained with Calcein-AM. Cells were stained with Lysobrite orange prior to imaging, TBP is detected in the 647 channel via Cy5 on the TBP. Scale bar corresponds to 20 μm.

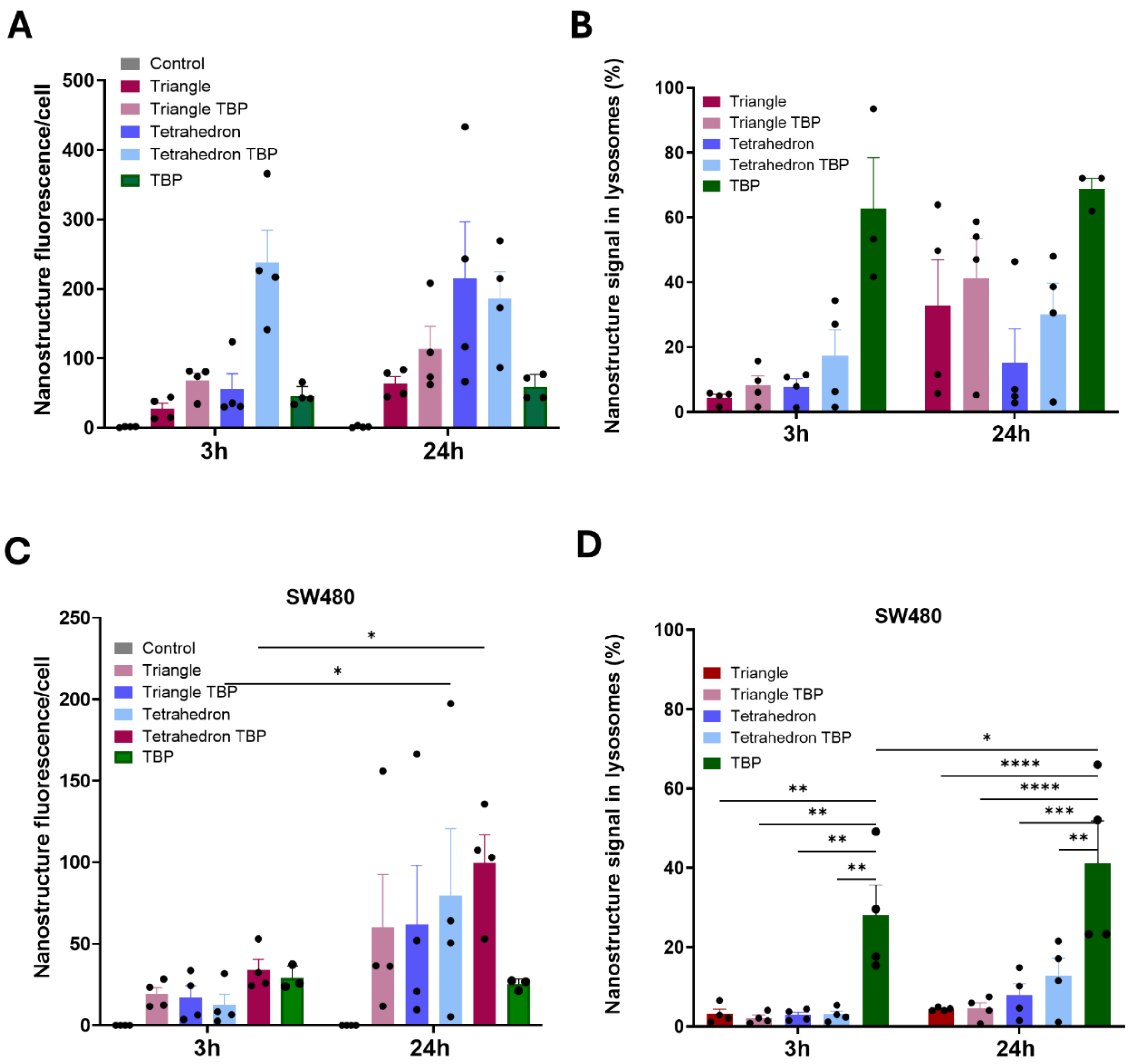

**Figure S9:** Endocytosis of TBP in HeLa Kyoto and SW80 cells, image quantification. A, C. Nanostructure internalisation represented as number of nanostructure fluorescence puncta per cell, four biological repeats, n>50 cells per biological repeat. B, D. Percentage of lysosomal colocalisation. Mean of technical replicates is plotted for the percentage of Alexa647 signal in lysosomes relative to the whole cell. Bars represent standard error of mean for biological replicates, and a two-way repeated measures ANOVA was used to determine significance (p<0.05=*, p<0.01=**, p<0.001=***, p<0.0001=****). Analysis performed via in-house developed CellProfiler pipeline.

**Table S1:** Descriptive statistics of Wnt signalling assay for HeLa Kyoto cell line.

| | Control (no treatment) | Triangle | Triangle TBP | Tetrahedron | Tetrahedron TBP | TBP only | IWR1 | G007-LK | No Wnt pathway activation |
|---|---|---|---|---|---|---|---|---|---|
| Number of values | 62 | 23 | 23 | 21 | 22 | 23 | 24 | 23 | 48 |
| Minimum | 39.0 | 34.0 | 20.0 | 38.0 | 32.0 | 38.0 | 26.0 | 13.0 | 2.06 |
| Maximum | 100 | 97.0 | 69.0 | 84.0 | 66.0 | 114 | 59.0 | 37.0 | 12.3 |
| Range | 61.0 | 63.0 | 49.0 | 46.0 | 34.0 | 76.0 | 33.0 | 24.0 | 10.2 |
| Mean | 67.3 | 60.1 | 46.0 | 60.3 | 46.6 | 73.5 | 42.9 | 25.0 | 6.52 |
| Std. Deviation | 16.4 | 19.6 | 12.6 | 13.6 | 9.00 | 18.7 | 10.0 | 7.93 | 2.47 |

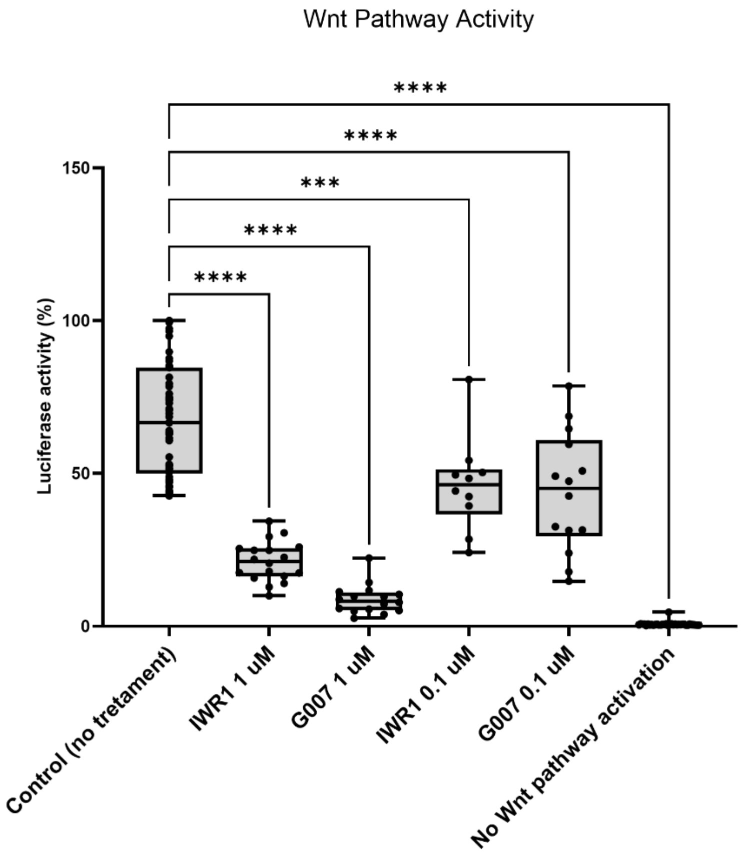

**Figure S10:** Potency of small molecule inhibitors IWR1, G007-LK in HEK-293T cells. Wnt signalling is very efficiently supressed at 1 uM final concentration of small molecule inhibitor, and at 0.1 μM concentration the effect is still significant, but more moderate. Statistical significance was determined through a one-way ANOVA ($p<0.05$=*, $p<0.01$=**, $p<0.0001$=****).

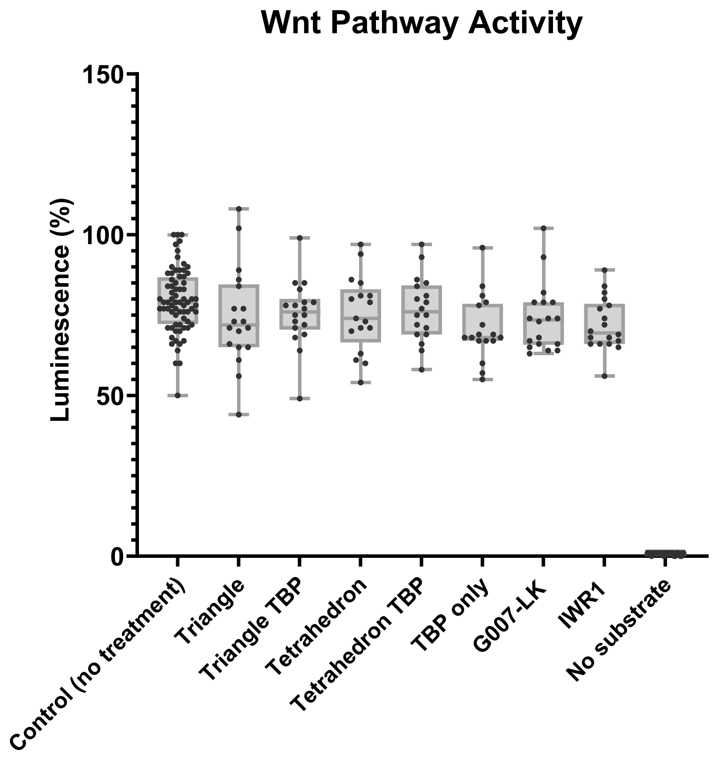

**Figure S11:** Wnt signalling assay results for SW480 cell line. after treatment with the nanostructures and small molecule TNKS inhibitors G007-LK and IWR1. Treatment concentrations are as follows: 3 nM triangle, triangle TBP (~100 nM effective TBP concentration), 50 nM tetrahedron, tetrahedron TBP (~100 nM effective TBP concentration), 100 nM free TBP, 1 μM IWR1, 1 μM G007-LK, incubation time 24h in serum-free media. Data are average of four biological repeats (>24 technical repeats). Statistical significance was determined through a one-way ANOVA (p<0.05=*, p<0.01=**, p<0.0001=****).

**Table S2:** Descriptive statistics of Wnt signalling assay for SW480 cell line

| | Control (no treatment) | Triangle | Triangle TBP | Tetrahedron | Tetrahedron TBP | TBP only | G007-LK | IWR1 | No substrate |
|---|---|---|---|---|---|---|---|---|---|
| Number of values | 72 | 18 | 18 | 17 | 18 | 17 | 18 | 18 | 36 |
| Minimum | 50.00 | 44.00 | 49.00 | 54.00 | 58.00 | 55.00 | 63.00 | 56.00 | 0.000 |
| Maximum | 100.0 | 108.0 | 99.00 | 97.00 | 97.00 | 96.00 | 102.0 | 89.00 | 1.000 |
| Range | 50.00 | 64.00 | 50.00 | 43.00 | 39.00 | 41.00 | 39.00 | 33.00 | 1.000 |
| Mean | 79.43 | 74.33 | 75.50 | 75.29 | 76.72 | 70.88 | 74.00 | 72.06 | 0.8333 |
| Std. Deviation | 9.919 | 15.49 | 10.29 | 11.85 | 10.08 | 10.20 | 10.57 | 8.207 | 0.3780 |
| Std. Error of Mean | 1.169 | 3.651 | 2.426 | 2.873 | 2.375 | 2.473 | 2.492 | 1.934 | 0.06299 |

**Table S3:** Multiple Comparisons of Wnt signalling assay for HeLa Kyoto cell line

| Comparisons | Mean diff. | Summary | Adjusted P Value |
|---|---|---|---|
| Control (no treatment) vs. Triangle | 7.187 | ns | 0.3911 |
| Control (no treatment) vs. Triangle TBP | 21.32 | **** | <0.0001 |
| Control (no treatment) vs. Tetrahedron | 6.941 | ns | 0.4893 |
| Control (no treatment) vs. Tetrahedron TBP | 20.68 | **** | <0.0001 |
| Control (no treatment) vs. TBP only | -6.248 | ns | 0.5893 |
| Control (no treatment) vs. IWR1 | 24.40 | **** | <0.0001 |
| Control (no treatment) vs. G007-LK | 42.27 | **** | <0.0001 |
| Control (no treatment) vs. No Wnt pathway activation | 60.75 | **** | <0.0001 |
| Triangle vs. Triangle TBP | 14.13 | * | 0.0103 |
| Triangle vs. Tetrahedron | -0.2464 | ns | >0.9999 |
| Triangle vs. Tetrahedron TBP | 13.50 | * | 0.0202 |
| Triangle vs. TBP only | -13.43 | * | 0.0188 |
| Triangle vs. IWR1 | 17.21 | *** | 0.0004 |
| Triangle vs. G007-LK | 35.09 | **** | <0.0001 |
| Triangle vs. No Wnt pathway activation | 53.56 | **** | <0.0001 |
| Triangle TBP vs. Tetrahedron | -14.38 | * | 0.0111 |
| Triangle TBP vs. Tetrahedron TBP | -0.6344 | ns | >0.9999 |
| Triangle TBP vs. TBP only | -27.57 | **** | <0.0001 |
| Triangle TBP vs. IWR1 | 3.082 | ns | 0.9968 |
| Triangle TBP vs. G007-LK | 20.96 | **** | <0.0001 |
| Triangle TBP vs. No Wnt pathway activation | 39.43 | **** | <0.0001 |
| Tetrahedron vs. Tetrahedron TBP | 13.74 | * | 0.0213 |
| Tetrahedron vs. TBP only | -13.19 | * | 0.0294 |
| Tetrahedron vs. IWR1 | 17.46 | *** | 0.0005 |
| Tetrahedron vs. G007-LK | 35.33 | **** | <0.0001 |
| Tetrahedron vs. No Wnt pathway activation | 53.81 | **** | <0.0001 |
| Tetrahedron TBP vs. TBP only | -26.93 | **** | <0.0001 |
| Tetrahedron TBP vs. IWR1 | 3.716 | ns | 0.9895 |
| Tetrahedron TBP vs. G007-LK | 21.59 | **** | <0.0001 |
| Tetrahedron TBP vs. No Wnt pathway activation | 40.07 | **** | <0.0001 |

| | | | |
|---|---|---|---|
| TBP only vs. IWR1 | 30.65 | **** | <0.0001 |
| TBP only vs. G007-LK | 48.52 | **** | <0.0001 |
| TBP only vs. No Wnt pathway activation | 67.00 | **** | <0.0001 |
| IWR1 vs. G007-LK | 17.88 | *** | 0.0002 |
| IWR1 vs. No Wnt pathway activation | 36.35 | **** | <0.0001 |
| G007-LK vs. No Wnt pathway activation | 18.48 | **** | <0.0001 |
